# RNase E resolves toxic condensates by counteracting phase separation of Type II RhlB helicases

**DOI:** 10.64898/2026.03.23.713837

**Authors:** Hausmann Stéphane, Johan Geiser, Oscar Vadas, Sylvain Guex-Crosier, Diego Gonzalez, Martina Valentini

## Abstract

In many Proteobacteria, the RNA helicase RhlB is a component of the RNA degradosome, a multi-protein complex involved in RNA processing and degradation. Within this complex, RhlB interacts with the scaffold endoribonuclease RNase E. In *Escherichia coli*, allosteric activation of RhlB by RNase E has defined the current paradigm for RhlB regulation. Here we identify a distinct clade of RhlB helicases, exemplified by *Pseudomonas aeruginosa* RhlB, which we designate Type II. Unlike Type I RhlB, Type II RhlB helicases contain an N-terminal intrinsically disordered region that drives RNA-dependent liquid-liquid phase separation and enhances RhlB activity. Biochemical, structural, and functional analyses show that *P. aeruginosa* RNase E binds RhlB through an interface distinct from that described in the *E. coli* model and, rather than stimulating activity, antagonizes RhlB phase separation. Excessive RhlB condensation impairs bacterial growth at low temperature, and RNase E–mediated control of RhlB condensation maintains growth under these conditions. Together, these findings reveal that conserved RNA degradosome components can engage in distinct regulatory interactions across species and identify condensate dissolution as a novel mechanism regulating RNA helicase activity.

## Introduction

RNA helicases are key enzymes that shape cellular RNA metabolism across all domains of life by remodeling RNA molecules and ribonucleoprotein complexes (1,2). By coupling ATP hydrolysis to the unwinding of double-stranded RNA regions and displacement of RNA-bound proteins, they assist processes such as transcription, translation, RNA decay, and ribosome biogenesis (3,4).

All RNA helicases share a conserved enzymatic core composed of two RecA-like domains connected by a flexible linker (5). These domains undergo conformational changes upon ATP and RNA binding and following ATP hydrolysis (6–8). By engaging primarily with the sugar–phosphate backbone of RNA, the enzymatic core allows RNA helicases to bind a broad range of RNA substrates without requiring primary sequence recognition (reviewed for example in 9,10).

Given their lack of sequence specificity, RNA helicase activity must be regulated to avoid unnecessary or deleterious RNA remodeling. This regulation is often achieved through auxiliary domains flanking the catalytic core, which can confer substrate specificity, modulate enzymatic activity, or facilitate interactions with regulatory cofactors (11,12). In many RNA helicases, auxiliary domains include intrinsically disordered regions that promote liquid–liquid phase separation (LLPS) in the presence of RNA, typically through weak, multivalent interactions (13–16). LLPS drives the formation of biomolecular condensates, dynamic assemblies that concentrate or sequester RNA, RNA helicases, and associated proteins, thereby generating localized biochemical environments (17,18). Through this membraneless compartmentalization, LLPS provides spatial control of RNA helicase activity and, more broadly, RNA metabolism. In eukaryotic cells, RNA helicase–driven LLPS underlies the assembly of membraneless organelles such as stress granules and processing bodies, which modulate gene expression in response to cellular demands (19–22).

In prokaryotes, LLPS of some RNA helicases has been shown, but its regulatory relevance remains poorly characterized. More broadly, only a few regulatory mechanisms controlling bacterial RNA helicase activity have been described (23–26). The allosteric control of RhlB by the endoribonuclease RNase E in *Escherichia coli* stands out as one of the best characterized examples. RhlB is a conserved DEAD-box RNA helicase present in many Proteobacteria, where it functions as part of the RNA degradosome, a multiprotein complex involved in RNA processing and degradation (27,28). In *E. coli*, RhlB exhibits low activity in isolation but is allosterically activated upon binding to RNase E, the endoribonuclease that scaffolds RNA degradosome assembly (27,29,30). The interaction occurs between the RNase E C-terminal extension (residues 698–762) and the RecA2 domain of RhlB (29,31–35). Stimulation of RhlB unwinding activity facilitates the degradation of structured RNAs and ribosome-free mRNAs (27,32,36). The functional enhancement through direct interaction with RNase E has served as the canonical model for RhlB regulation in bacteria.

Our laboratory recently identified RhlB (PA3861) as a core component of the RNA degradosome in the bacterial pathogen *Pseudomonas aeruginosa* (37). Unlike in *E. coli*, this interaction involves a distinct RNase E binding site (residues 757–776, corresponding to the AR1 <u>s</u>hort <u>li</u>near <u>m</u>otif or SLiM), prompting us to investigate how *P. aeruginosa* RhlB is regulated. Here, we show that *P. aeruginosa* RhlB defines a phylogenetically and functionally distinct clade of RhlB helicases, named Type II RhlB. *P. aeruginosa* RhlB undergoes RNA-dependent LLPS driven by an intrinsically disordered N-terminal extension, which also enhances the protein catalytic activity. RNase E binds within the RecA1 domain of *P. aeruginosa* RhlB, rather than the RecA2 domain as in *E. coli*. Instead of stimulating RhlB activity, RNase E suppresses *P. aeruginosa* RhlB phase separation. This regulatory mechanism is critical for bacterial growth at low temperature, where uncontrolled RhlB condensation becomes detrimental. This study reveals phase separation as a regulatory layer controlling Type II RhlB RNA helicase activity and establishes RNase E as a central regulator of this process. The composition of the RNA degradosome is known to vary substantially across bacterial species, reflecting distinct combinations of ribonucleases, helicases, and accessory factors (38,39). Here, we show that the RNA degradosome plasticity arises not only from compositional variation but also from divergent interaction modes among conserved RNA degradosome components.

## Results

### PaRhlB forms a distinct RhlB clade with an intrinsically disordered N-terminal extension driving LLPS

Genome analysis predicted that *P. aeruginosa* RhlB (PaRhlB) contains an N-terminal extension (NTE) absent in the *E. coli* homolog (EcRhlB) (40). We confirmed the NTE presence by performing a mass spectrometry analysis of chromosomally C-terminally tagged PaRhlB–mRuby2 purified from *P. aeruginosa* and detecting peptides within the NTE (Table S1).

To investigate the evolutionary distribution of the NTE among RhlB homologs, we performed a phylogenetic analysis (Fig. 1A and Fig. S1). This revealed that RhlB proteins separate into two main clades according to the presence of the NTE. The more widespread Type I clade includes EcRhlB and homologs lacking an NTE, whereas the Type II clade includes PaRhlB and other Pseudomonads RhlB proteins with NTEs (Fig. S1-S2). In the analysis, we included *C. crescentus* RhlB (CC1847), which did not cluster with RhlB homologs, consistent with its previous classification as RhlE-like (41).

**Figure 1:**
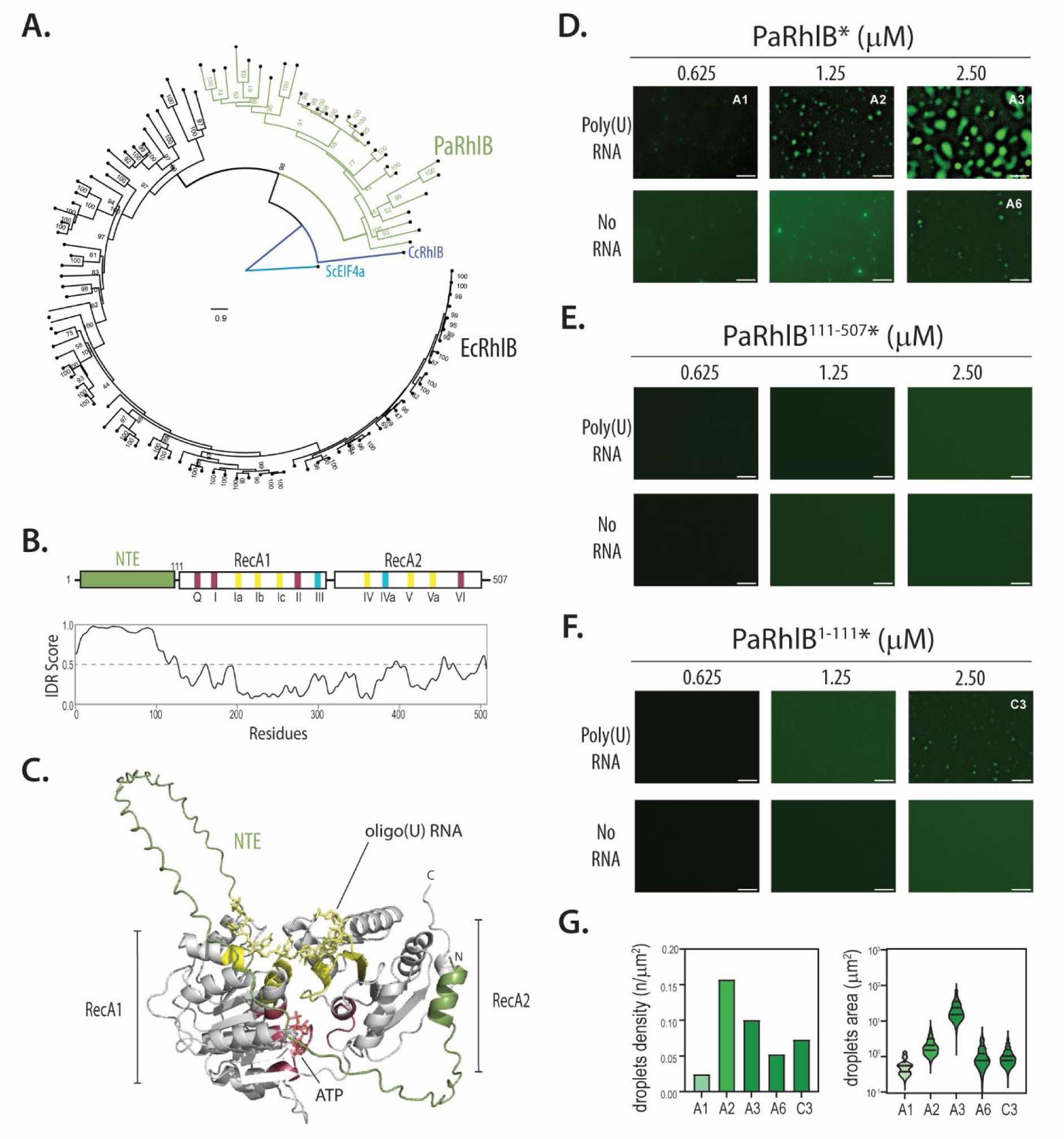
The NTE stimulates *Pa*RhlB LLPS. (**A**) Phylogenetic tree of RhlB homologs. Green branch refers to the Type II RhlB branch, including PaRhlB. EcRhlB and CcRhlB indicate *E. coli* and *C. crescentus* proteins, respectively. The extended tree is shown in Fig. S1. *Saccharomyces cerevisiae* EIF4a RNA helicase (ScEIF4a) was used as outgroup. (**B**) Domain organization of PaRhlB and prediction of intrinsically disordered regions using AIUPred (8). The conserved RNA helicase motifs (Q–VI) are indicated. The N-terminal extension (NTE) is shown in green; motifs involved in ATP binding and hydrolysis are shown in red; those mediating RNA binding are in yellow; and motifs contributing to intramolecular interactions between the two RecA-like domains are in blue. (**C**) Alphafold3 prediction of *Pa*RhlB-RNA in the presence of ATP (red)(61). Motifs colors as in panel B. Olig(U) RNA in yellow and ATP in red. **(D–F)** In vitro droplet formation of PaRhlB or its variants (final concentrations: 0.625 µM, 1.25 µM, or 2.5 µM) in the presence of 250 ng/mL poly(U) RNA (top row) or in its absence (bottom row). Note that * indicates that protein mixtures contained the indicated RhlB variant together with its msfGFP-tagged version at a 100:1 molar ratio (unlabeled:msfGFP-labeled). (**G**) Quantification of droplets density (left panel) and droplets area (right panel) of images A1, A2, A3, A6 and. All LLPS assays were performed in three independent experiments; representative images are shown.

Structural predictions indicate that the NTE is intrinsically disordered (Fig. 1B-C). Intrinsically disordered regions (IDRs) are known to promote liquid–liquid phase separation (LLPS) (16,42). To examine phase separation behavior of PaRhlB, N-terminal His_10_–Smt3 tagged PaRhlB, PaRhlB^111–507^ (lacking the NTE) and the isolated NTE (PaRhlB^1–111^) were purified by nickel–agarose affinity chromatography (Fig. S3). The corresponding msfGFP–tagged proteins were purified in parallel, and in vitro LLPS assays were performed by mixing the msfGFP-tagged and untagged versions at a 1:100 ratio, respectively (see Materials and Methods). PaRhlB formed numerous visible droplets at a minimum concentration of 1.25 µM in an RNA-dependent manner (Fig. 1D). Area occupied by droplets increased proportional to protein concentration, while a decrease in droplet density was observed when comparing 2.5 and 1.25 µM, suggesting droplets fusion (Fig. 1G). FRAP analysis of RhlB droplets revealed a mobile fraction of 78 ± 2 % and a recovery half time of 15 ± 1 s (Fig. S4), consistent with a dynamic, liquid-like condensate. In contrast, the PaRhlB^111–507^ truncation failed to undergo phase separation in vitro, even at 2.5 µM, indicating that the NTE is required for efficient LLPS (Fig. 1E). Of note, although the His_10_–Smt3 tag facilitated droplet formation, it did not influence the observed differences in protein phase separation (Fig. S5A). Interestingly, the purified His_10_–Smt3 tagged NTE fragment (PaRhlB^1-111^) was able to phase separate in the presence of RNA, although less efficiently than RhlB (Fig. 1F), suggesting that the RecA-like core domain also contributes to droplet formation. We were unable to assess the untagged NTE because removal of the His_10_–Smt3 tag resulted in aggregate formation, suggesting that the NTE must remain linked to a folded domain to remain soluble.

### The PaRhlB NTE enhances core helicase activities

We next examined the role of the NTE in PaRhlB biochemical activity, beginning with its RNA-dependent ATP hydrolysis. Full-length PaRhlB catalyzed the release of inorganic phosphate from ATP only in the presence of poly(U) RNA (≥150 ribonucleotides, heterogeneous in size; Fig. 2A). The ATP hydrolysis rate, determined from the linear range of activity (0–0.125 µM RhlB), corresponded to an estimated turnover number of ∼98 min⁻¹. In contrast, the NTE-truncated variant PaRhlB^111–507^ retained only ∼10% of the wild-type ATPase activity, indicating that the NTE is critical for RNA-dependent catalysis (Fig. 2A). This residual activity was still dependent on poly(U) RNA (Fig. 2A). Mutation of the Walker A motif within the catalytic domain of the protein (PaRhlB^K169A^) reduced activity to <1% of wild type (Fig. S5B), confirming that the ATPase activity is intrinsic to the recombinant protein.

**Figure 2:**
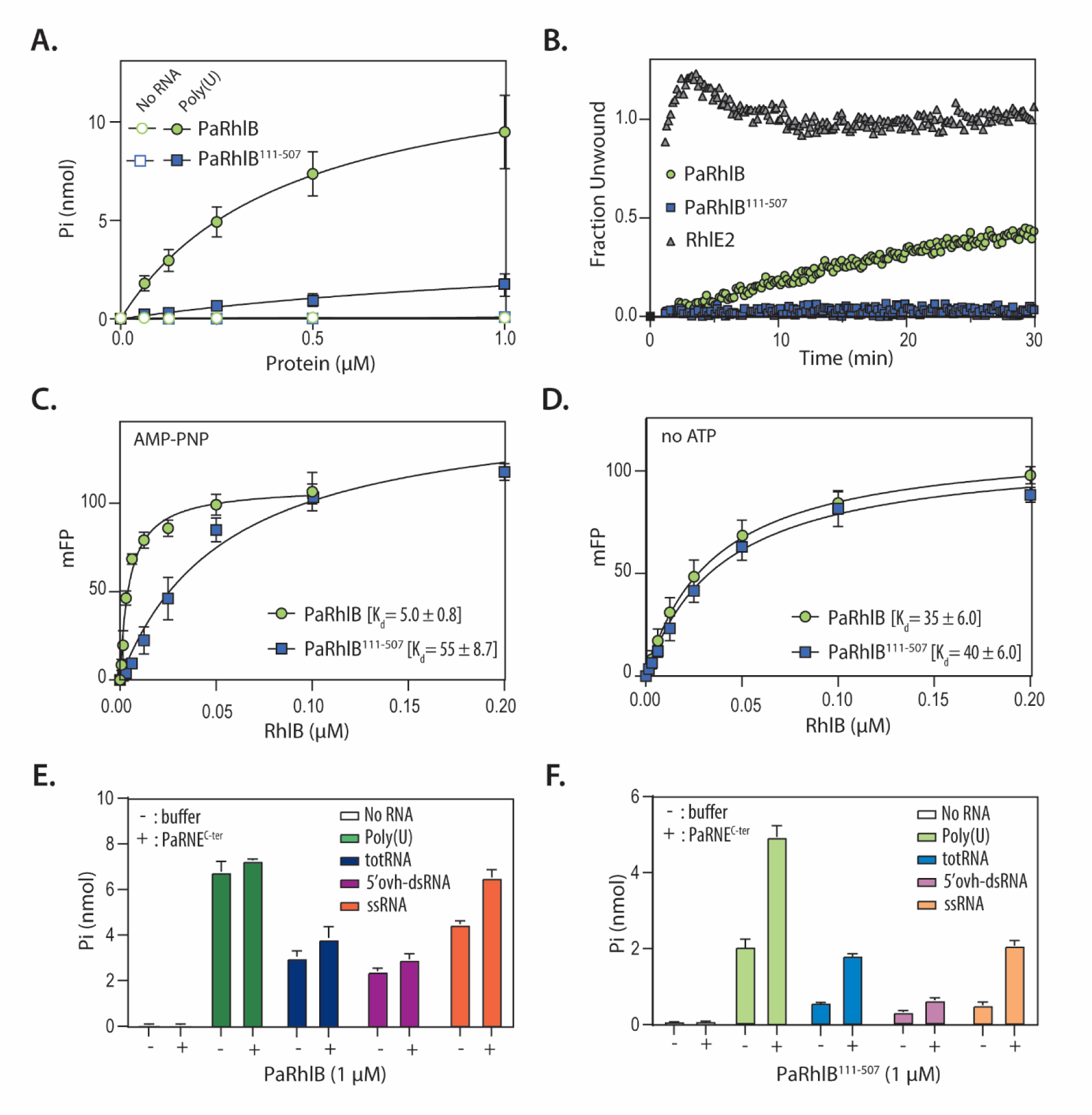
The NTE of *Pa*RhlB promotes RhlB activities and ATP-dependent RNA binding. (**A**) RNA-dependent ATPase activity of full length PaRhlB (RhlB^1-507^) and the NTE deleted variant RhlB^111-507^. Pi release is plotted as a function of input protein. Data are the average of three independent experiments with error bars representing standard deviation (SD). (**B**) RNA unwinding activity of RhlB^1-507^ and RhlB^111-507^. The extent of RNA unwound is plotted as a function of time. (**C**) ATP-dependent RNA binding of RhlB^1-507^ and RhlB^111-507^. mFP is plotted as of a function of protein concentration. Data are the average of six independent experiments with error bars representing SD. (**D**) ATP-independent RNA binding of RhlB^1-507^ and RhlB^111-507^. Experiments were performed as described in (C), except that AMP-PNP was omitted. (**E-F**) ATPase activity of 1 µM RhlB^1-507^ (E) and RhlB^111-507^ (F) in the absence (–, buffer) or presence (+) of 1 µM PaRNase E^C-ter^ (RNE ^C-ter^) with the indicated RNA substrates (none, Poly(U), totRNA, 5′ovh-dsRNA, and ssRNA]. Mean ± SD of three independent experiments.

RNA unwinding activity was assessed using a real time fluorescence-based helicase assay (see Materials and Methods). Briefly, a 5′ovh_dsRNA_26/14_ substrate (consisting of a 5′-Cy5–labelled 14-mer RNA annealed to a 26-mer RNA carrying a 3′ quencher) was used, and unwinding was monitored by increased fluorescence in the presence of ATP and Mg²⁺ over 30 min due to strand separation. *P. aeruginosa* RhlE2 served as positive control (26). Under these conditions, 50 nM of PaRhlB unwound ∼45% of the duplex RNA (Fig. 2B). This activity, although weak when compared to RhlE2, was absent in the PaRhlB^K169A^ mutant and strictly ATP-dependent (Fig. S6A-B). Substrate specificity tests revealed that PaRhlB cannot unwind blunt-ended RNA duplexes or duplexes with a 3′ overhang (Fig. S6C), like EcRhlB (29,34,43). In agreement with its poor ATPase activity, PaRhlB^111–507^ showed only <1% of full-length PaRhlB helicase activity (Fig. 2B).

To determine RNA binding affinity, we performed fluorescence polarization assays using a 5-carboxyfluorescein–labeled 5′ overhang dsRNA_26/14_. Since ATP can influence RNA binding affinity of DEAD-box RNA helicases (3,44), we performed the assays in the presence or absence of the non-hydrolysable ATP analogue adenosine-5′-(β,γ-imido)triphosphate (AMP-PNP) and Mg²⁺. In the presence of AMP-PNP, full-length PaRhlB bound RNA with ∼10-fold higher affinity than PaRhlB^111-507^ (Kd = 5 nM vs. 55 nM) (Fig. 2C). In the absence of AMP-PNP, the RNA-binding affinity of full-length PaRhlB decreased ∼7-fold (Kd = 35 nM), whereas PaRhlB^111-507^ showed only a minimal change (Kd = 40 nM vs. 55 nM) (Fig. 2D). These results indicate that the contribution of the NTE to RNA binding depends on ATP binding to the helicase core.

As the proteins carry an N-terminal His_10_–Smt3 tag, we verified that the tag did not affect the RNA-dependent ATPase activity, RNA unwinding, nor RNA binding (Fig. S5B-D).

### RNase E stimulation of PaRhlB activity is hindered by the NTE

Using direct binding assays and a bacterial two-hybrid (BTH) system, we have previously demonstrated that PaRhlB interacts with the AR1 SLiM of *P. aeruginosa* RNase E (PaRNase E) (37). Direct binding assays between RhlB^111-507^ and RNase E C-terminal scaffolding region (PaRNase E^C-ter^, residues 572-1057) revealed that the NTE is not required in the interaction (Fig. S7).

Guided by the EcRhlB model, we next tested whether PaRNase E could stimulate the ATPase activity of PaRhlB. In the presence of equimolar PaRNase E^C-ter^, full-length PaRhlB showed no significant stimulation by RNase E in the presence of poly(U) RNA (Fig. 2E and S8A). In contrast, PaRhlB^111–507^ ATPase activity increased consistently by ∼2-fold across all protein concentrations. Doubling the concentration of PaRNase E did not result in further stimulation (Fig. S8A).

The degree of activation of EcRhlB by EcRNase E changes using different RNA substrates (45), we therefore measured ATPase activity of either full-length PaRhlB or PaRhlB^111–507^ using a range of RNA substrates: *P. aeruginosa* total RNA (totRNA), a 5′ overhang dsRNA_26/14_ (5′ovh-dsRNA), and a 20-mer single-stranded RNA (ssRNA). Negative controls included reactions without RNA (buffer). Again, PaRNase E^C-ter^ did not appreciably stimulate full-length PaRhlB (maximum ∼1.2-fold increase for ssRNA), whereas PaRhlB^111–507^ was stimulated between 2- and 4-fold depending on the RNA substrate (Fig. 2E-F). Taken together, these results show that PaRNase E stimulates ATPase activity only in the truncated PaRhlB^111–507^. The ATP-dependent contribution of the NTE to PaRhlB RNA binding, observed earlier (Fig. 2C), may therefore render additional stimulation by RNase E unnecessary in the full-length protein.

### Identification of PaRhlB sites mediating PaRNase E binding by HDX-MS

To map the region(s) of PaRhlB involved in PaRNase E interaction in an unbiased manner, we performed hydrogen–deuterium exchange coupled to mass spectrometry (HDX–MS). PaRhlB, either alone or in complex with PaRNase E^C-ter^ was incubated in deuterated buffer for 3 s, 30 s, or 300 s before quenching the reaction at low pH and temperature (Table S2). Deuterium incorporation was quantified by pepsin digestion followed by UPLC–MS analysis. HDX rates for apo PaRhlB and PaRhlB in the presence of PaRNase E^C-ter^ were calculated for each peptide. In total, 149 PaRhlB peptides were identified and characterized under all conditions, providing a sequence coverage of 75% (Fig. S9 and Tables S3a-c).

Overall, two main regions within the RecA1 domain of PaRhlB exhibited a significantly reduced HDX rate (p-value< 0.01) in the presence of PaRNase E^C-ter^. These regions span residues 182–197, located upstream of motif Ia, and residues 234–245 upstream of motif Ib (Fig. 3A-C). HDX–MS findings align with BTH data, which also identified RecA1 as the primary PaRNase E–binding domain (Fig. 3D). EcRNase E was previously reported to interact primarily with the RecA2 domain of EcRhlB, with little contribution from RecA1 (33,34). These results suggest that although the interaction between RNase E and RhlB is conserved, the specific domain interface differs between PaRhlB and EcRhlB. To explore this further, we tested heterologous interactions using the BTH system (Fig. 4A). PaRNase E interacted with both PaRhlB and EcRhlB, whereas EcRNase E bound EcRhlB and the truncated variant PaRhlB^111-507^, but not the full-length PaRhlB, indicating that the NTE hinders the interaction. BTH assays further indicated that EcRNase E interacts with the RecA1 domain of PaRhlB, rather than with RecA2.

**Figure 3:**
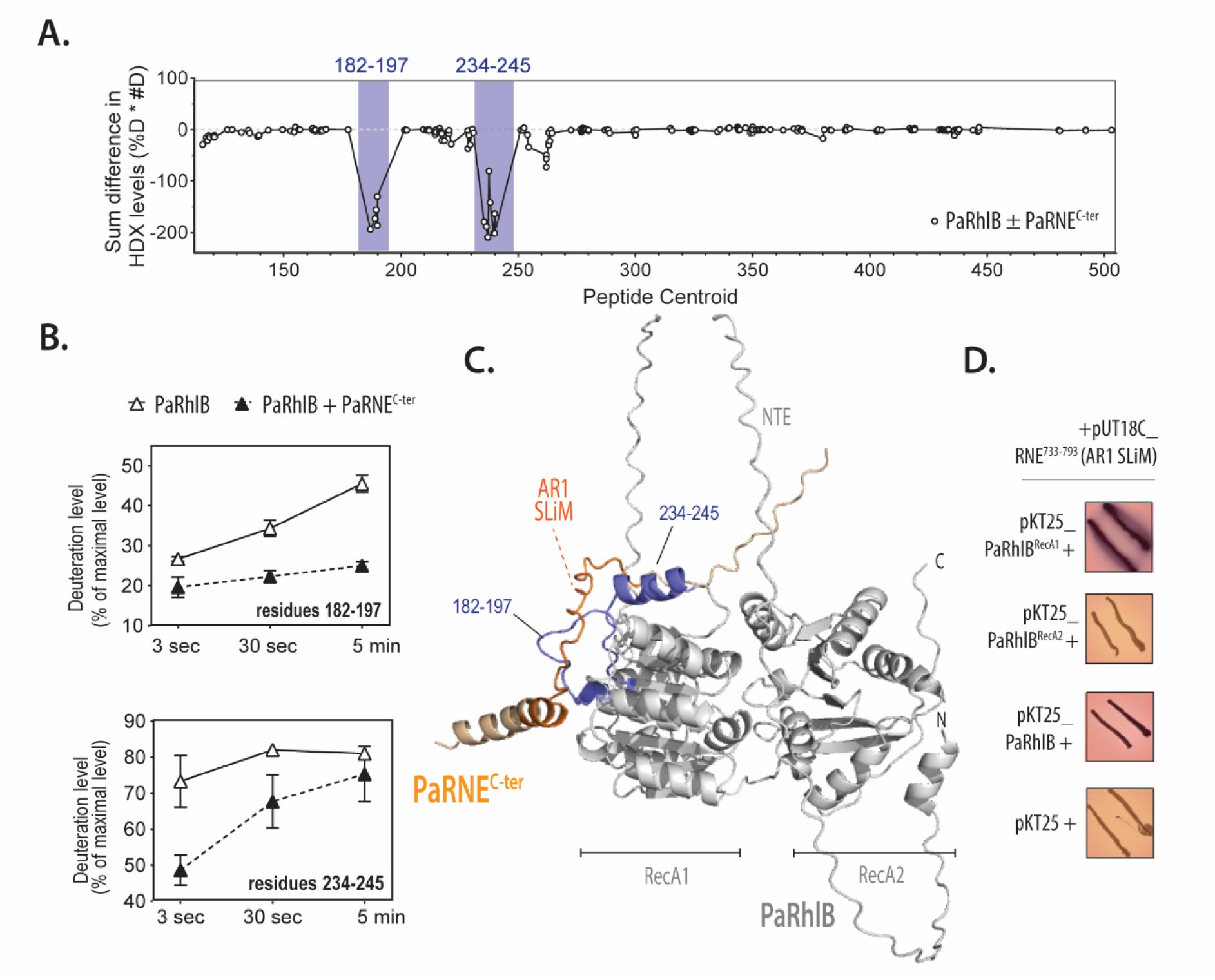
Identification of RNase E interaction regions on RhlB by HDX-MS. (**A**) Linear plot showing changes in deuterium uptake (HDX) in RhlB upon binding to RNase E ^C-ter^ (RNE ^C-ter^: residues 572-1057), plotted as a function of the central residue number of each peptide (x-axis). (**B**) Deuterium uptake kinetics (% deuteration ± standard deviation) over time for selected regions of RhlB. The corresponding residue numbers are indicated within each graph. (**C**) Structural mapping of RNase E interaction sites on the PaRhlB AlphaFold3 predicted structure (61). Peptides showing significant HDX changes in the presence of RNE ^C-ter^ are colored in blue. The RNase E segment is shown in orange. PyMOL was used for structural visualization (62). (**D**) Bacterial two-hybrid (BTH) assay mapping the PaRhlB (RecA1, RecA2 domains or full length) binding to the AR1 SLiM of PaRNase E^733-793^. BTH assays were performed in three independent experiments; representative images are shown.

**Figure 4:**
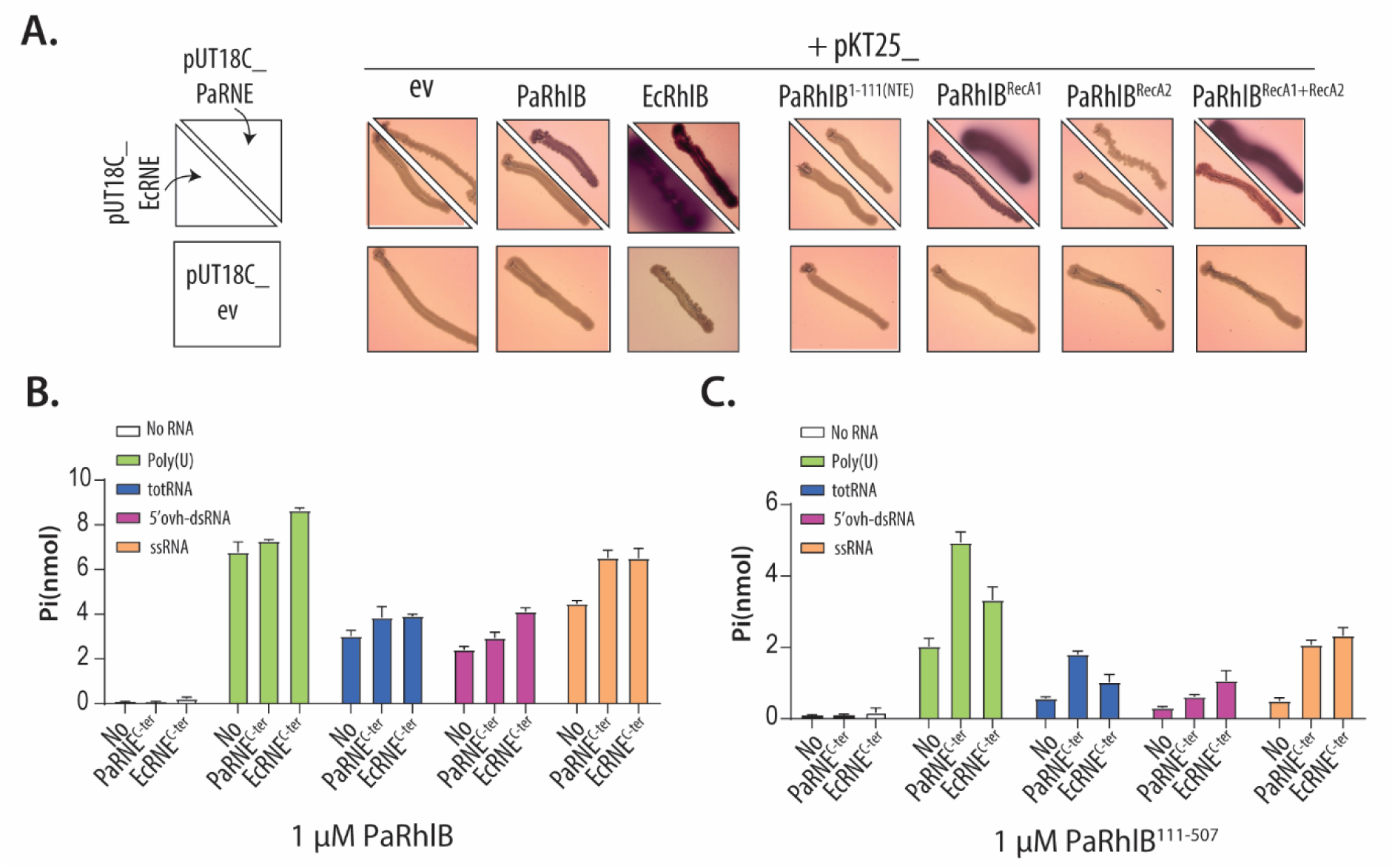
Heterologous interactions and stimulation of PaRhlB ATPase activity. (**A**) Detection of heterologous interactions using the bacterial two-hybrid (BTH) system. Reconstitution of adenylate cyclase in *E. coli* BTH101 was monitored by the appearance of red colonies on McConkey agar supplemented with 1% maltose, 0.5 mM IPTG, 100 µg/ml ampicillin, and 50 µg/ml chloramphenicol. Each strain co-expressed one pUT18C construct (pUT18C–PaRNase E, pUT18–EcRNase E, or empty vector) together with one pKT25 construct (indicated above each image). Red coloration indicates interaction-dependent cAMP production driving maltose fermentation and medium acidification. (**B–C**) ATPase activity of 1 µM PaRhlB (**B**) or PaRhlB^111-507^ (**C**) measured in the absence (No) or presence of 1 µM PaRNE^C-ter^ (RNase E^572-1057^) or 1 µM EcRNE^C-ter^ (EcRNase E^588-1061^), using the indicated RNA substrates (none, poly(U), totRNA, 5′ovh-dsRNA, or ssRNA). Data represent the mean ± SD of three independent experiments. See Fig S8 for extended data.

Functionally, EcRNase E^588-1061^ (EcRNase E^C-ter^) stimulated the ATPase activity of PaRhlB^111-507^ by approximately 2- to 7-fold, depending on the RNA substrate tested, while no significant activation was observed for full-length PaRhlB (Fig. 4B-C). This suggests that both binding modes can promote RhlB activation, albeit with different efficiencies. Under the same assay conditions, EcRNase E^C-ter^ enhanced the ATPase activity of EcRhlB by 2.3- to 14-fold across the RNA substrates tested, whereas stimulation ranged from 5- to 25-fold in the buffer system described by Zetzsche *et al.* (2023) (Fig. S8B-C, 50 mM KGlu vs 100 mM KCl, respectively), consistent with their finding (45). These results also validate our observations with PaRhlB^111–507^, demonstrating that stimulation is not an artifact of assay conditions.

### PaRNase E impairs PaRhlB phase separation

In *C. crescentus*, RNase E undergoes LLPS to form BR-bodies, biomolecular condensates that promote mRNA degradation and contain RhlB as well as other RNA degradosome components (46). We therefore tested whether PaRhlB LLPS could be influenced by PaRNase E. Two protocols were used. In the first, PaRhlB (PaRhlB-msfGFP:PaRhlB, 1:100) was pre-incubated with poly(U) RNA for 60 min before adding PaRNase E^C-ter^ (residues 572-1057) and incubating for another 60 min (Fig. 5A). In the second, the same mixture was co-incubated with PaRNase E^C-ter^ and poly(U) RNA for 60 min (Fig. 6A). PaRhlB droplet area and partitioning ratio were quantified. Unexpectedly, adding PaRNase E^C-ter^ to pre-formed PaRhlB droplets caused complete dissolution, and no PaRhlB droplet formation occurred in its presence (Fig. 5B and 6B, respectively).

**Figure 5:**
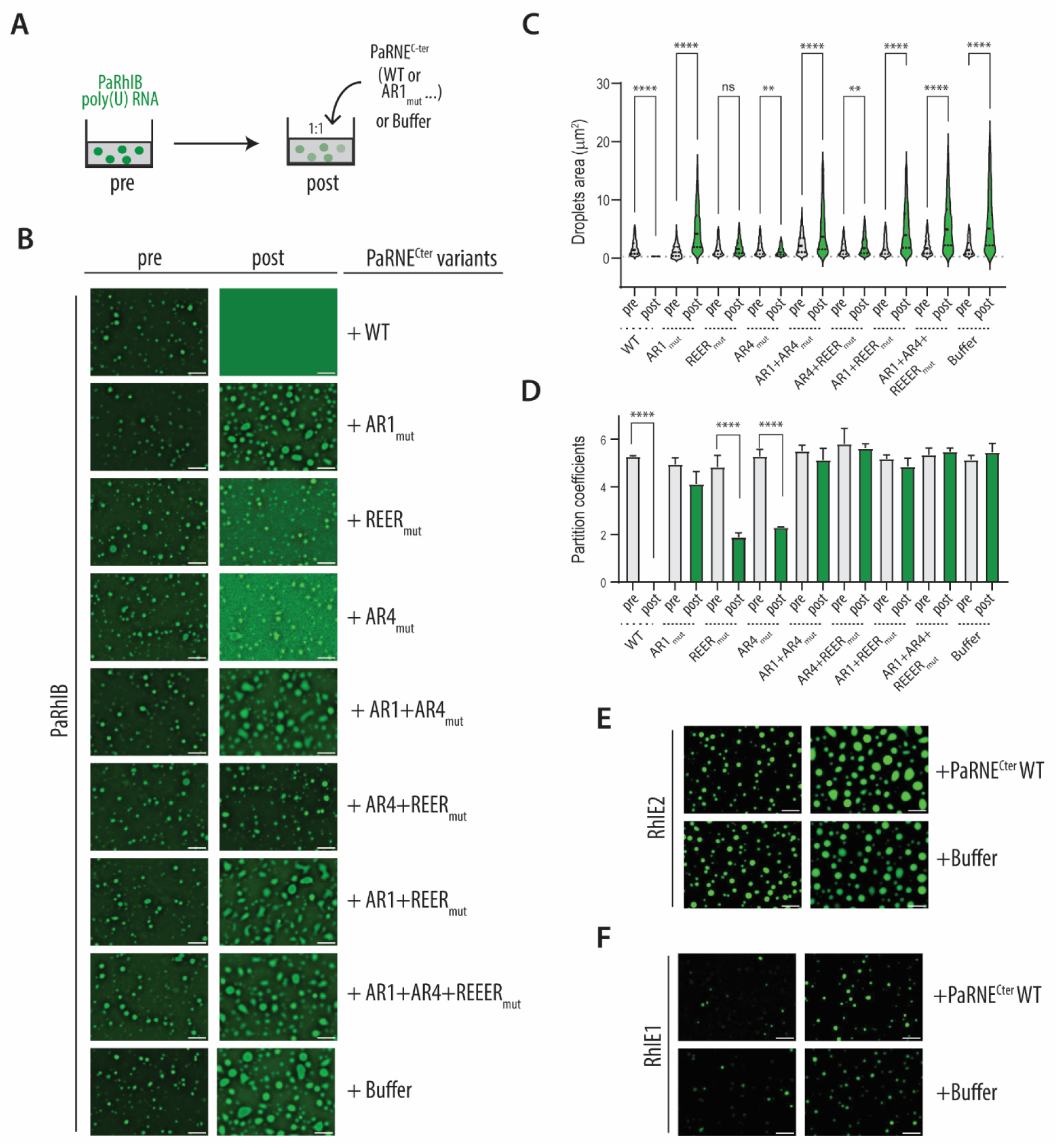
*P. aeruginosa* RNase E dissolves *Pa*RhlB LLPS in an AR1- and RNA-dependent manner. (**A**) Schematic workflow of the RhlB droplet dissolution assay using RNase E ^C-ter^ (RNE ^C-ter^: residues 572-1057) WT or its mutant variants. (**B**) In vitro droplet formation of 2.5 µM RhlB in the presence of 250 ng/mL poly(U) RNA (first column) and following addition of 2.5 µM RNE ^C-ter^ or its mutant variants. In the first column, images were captured 60 minutes after induction of phase separation with poly(U). In the second column, images were taken 60 minutes after the addition of RNE ^C-ter^ WT, its variants, or buffer (control). (**C**) Quantification of droplet area from three independent replicates. (**D**) Quantification of the partition coefficient from three independent replicates (**E–F**) In vitro droplet formation of 2.5 µM RhlE2 (**E**) or RhlE1 (**F**) in the presence of 250 ng/µl poly(U) RNA (left column) and upon addition of 2.5 µM RNE ^C-ter^ (buffer used as a negative control).

**Figure 6:**
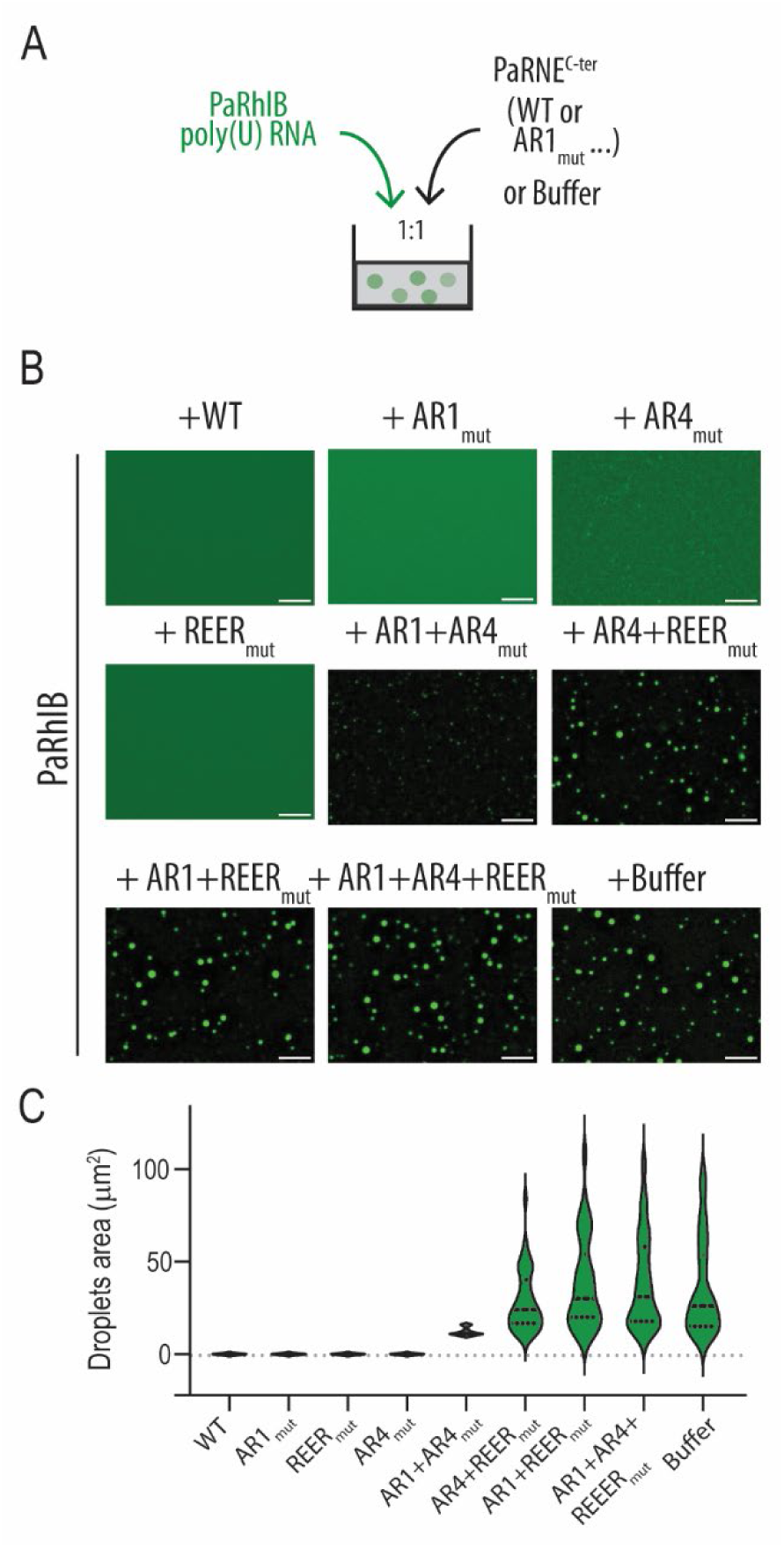
*Pa*RNase E impairs *Pa*RhlB LLPS in an AR1- and RNA-dependent manner. (**A**) Schematic workflow of the RhlB droplet formation assay using RNase E ^C-ter^ (RNE ^C-ter^: residues 572-1057) WT or its mutant variants. (**B**) Fluorescent microscopy images of 2.5 µM RhlB droplets in the presence of 250 ng/mL poly(U) RNA and RNE ^C-ter^ WT or mutated variants. (**C**) Quantification of area occupied by droplets. LLPS formation assays were performed in three independent experiments; representative images are shown.

We next investigated whether the SLiMs in PaRNase E^C-ter^ that mediate PaRhlB and RNA binding contribute to its inhibitory effect. The AR1 SLiM interacts with both PaRhlB and RNA, whereas the AR4 and REER SLiMs bind RNA only (37). To probe their roles, we used previously generated PaRNase E^C-ter^ mutated variants carrying alanine-replacements targeting each SLiM: AR1mut (12 Arg→Ala), AR4mut (8 Arg + 2 Gln→Ala), and REERmut (41 Arg→Ala), alone or in combination (37). Mutation of AR1 abolished droplet dissolution: PaRhlB droplets were not dispersed but continued to fuse and increase in size over time (Fig. 5B–C). On the contrary, PaRhlB droplet formation was still impaired by AR1mut similarly to the WT (Fig 6B-C). AR4mut and REERmut retained partial dissolution activity, as indicated by the absence of droplet area increase between pre- and post-treatment and by reduced partition coefficients, and they inhibited droplet formation to a similar extent as the WT and AR1mut (Fig. 5C–D and Fig. 6C). Combining two or three SLiM mutations produced additive effects, and the triple mutant fully lost any inhibitory activity, behaving like the buffer control. Together, these results demonstrate that both RhlB binding (via AR1) and RNA binding (via AR1, AR4, and REER) are required for PaRNase E–mediated inhibition of PaRhlB droplets, with AR1 playing the dominant role in dissolution of already formed droplets.

We also examined PaRNase E^C-ter^ effect on *P. aeruginosa* RhlE2 and RhlE1 LLPS (26). Under identical conditions, RhlE2 and RhlE1 droplets remained unaffected (Fig. 5E-F), indicating that PaRNase E^C-ter^ selectively targets PaRhlB. Finally, PaRNase E^C-ter^ efficiently dissolved PaRhlB droplets formed with poly(U), MS2, Pa totRNA, whereas EcRNase E^C-ter^ (residues 588-1061) failed to do so, consistent with its lack of interaction with PaRhlB (Fig. S10). Of interest, PaRhlB droplet size varied with the RNA substrate, being smallest with Pa totRNA.

### *Pa*RNase E counteracts *Pa*RhlB NTE-dependent cold-induced toxicity

Phenotypic analyses of *P. aeruginosa* Δ*rhlB* mutant revealed a mild defect in colony formation at 16 °C compared to the wild type, whereas no growth difference appeared at 37 °C, indicating a cold-sensitive phenotype.

Unexpectedly, RhlB overexpression caused a strong growth defect at low temperature in both wild-type and Δ*rhlB* backgrounds. This effect was independent of PaRhlB catalytic activity, as overexpression of the ATPase-deficient mutant PaRhlB^K169A^ produced a growth defect comparable to that observed with the wild-type protein. In contrast, the truncated variant PaRhlB^111–507^ (missing the PaRhlB NTE) had no effect on growth (Fig. 7A). A similar phenotype was observed in liquid cultures at 18 °C (Fig. 7B), and comparable protein expression levels were confirmed by Western blot (Fig. S11A). Together, these results indicate that PaRhlB supports growth at low temperature, but its overexpression is toxic in an NTE-dependent manner, suggesting a deleterious effect of uncontrolled phase separation.

**Figure 7:**
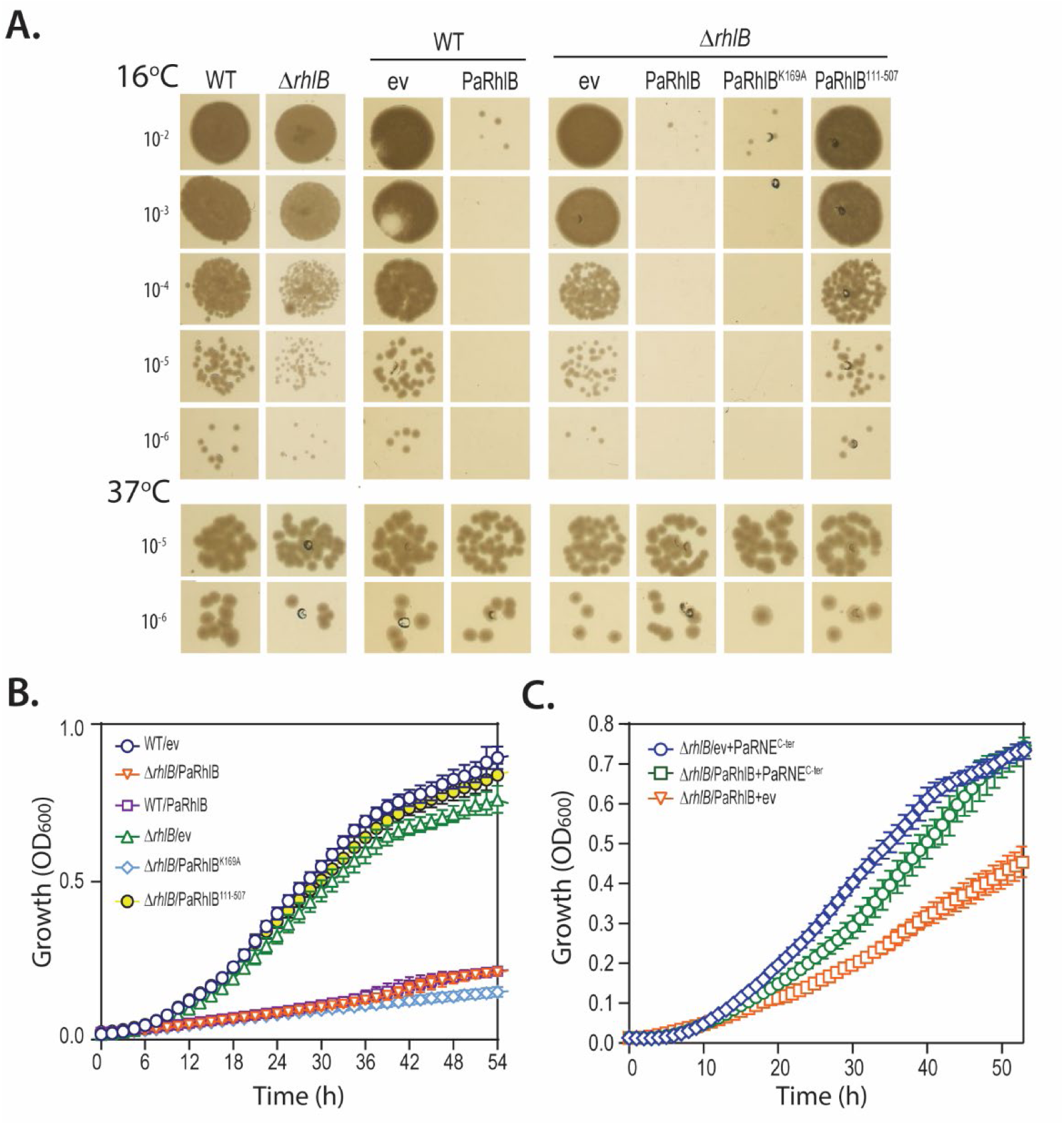
PaRhlB influences *P. aeruginosa* growth on cold and droplets cold growth. (**A**) Growth of *P. aeruginosa* wild-type (WT) or Δ*rhlB* mutant strains on nutrient agar (NA) plates at 16°C or 37°C, either alone or carrying plasmids for overexpression of RhlB, RhlB^K169A^ or RhlB^111-507^ variants. Ev: empty vector control. The vector used to overexpress RhlB carried the strong PEM7 constitutive promoter (63). Experiments were performed in biological independent triplicates; a representative plate is shown. Protein expression levels were verified by western blotting (see Fig. S10A). (**B**) Growth of the same strains shown in panel A in nutrient yeast broth (NYB) at 18°C. (**C**) Growth curves of Δ*rhlB* mutant strains co-expressing RhlB (IPTG, 0.3 mM) and/or RNase E ^C-ter^ (RNE ^C-ter^: residues 572-1057, arabinose, 0.03%) at 18 °C, controls strains carry empty vectors (ev). Each value is the average of three different cultures ± the standard deviation. In some instances, the standard deviation bars are smaller than the symbols used.

Given that PaRNase E^C-ter^ (residues 572–1057) dissolves or impairs PaRhlB droplet formation *in vitro*, we reasoned that co-expression of PaRNase E^C-ter^ might alleviate PaRhlB toxicity in vivo. To modulate RhlB expression levels, *rhlB* was placed under the control of an IPTG-inducible promoter. Reducing PaRhlB levels resulted in a milder but still significant growth defect that increased with IPTG concentration, consistent with a dose-dependent effect (Fig. S11B). PaRNaseE^C-ter^ was then expressed from a second vector under the control of an arabinose-inducible promoter. In agreement with our hypothesis, PaRNAse E^C-ter^ rescued the growth defect caused by PaRhlB induction, restoring growth to the levels observed in the absence of RhlB (Fig. 7C). Altogether, these results indicate that PaRhlB LLPS is deleterious for bacterial growth at low temperature and that PaRNase E can counteract this effect by dissolving droplets or preventing their formation.

## Discussion

Our study identifies *P. aeruginosa* RhlB (PaRhlB) as the representative of a phylogenetically and functionally distinct subgroup of RhlB helicases (Type II), which differs from the canonical Type I group exemplified by *E. coli* RhlB (EcRhlB). A hallmark of Type II RhlBs is the presence of an intrinsically disordered N-terminal extension (NTE). We show that, in PaRhlB, this NTE promotes liquid–liquid phase separation (LLPS) and enhances the activity of the core RecA-like helicase domain, increasing ATP-stimulated RNA binding, RNA-dependent ATP hydrolysis, and RNA unwinding. In contrast, EcRhlB lacks an NTE and relies on interaction with RNase E, the central component and scaffold of the RNA degradosome, for catalytic activation (34,35,45).

A second key insight from this work is that, although RNase E remains a conserved partner of RhlB across diverse bacteria, the mode of interaction between the two proteins differs. In *P. aeruginosa*, we show that the interaction is mediated by the RecA1 domain of PaRhlB and the RNase E AR1 SLiM, rather than by the canonical interface of *E. coli*, where the RhlB RecA2 domain engages the RNase E region adjacent to the RBD–AR2 SLiMs (amino acids 698–762) (29,31–35).

Functionally, PaRNase E does not stimulate PaRhlB catalytic activity (as observed in *E. coli*) but instead inhibits PaRhlB LLPS. This inhibition requires the PaRNase E AR1, AR4, and REER SLiMs, indicating that multivalent contacts with both PaRhlB and RNA contribute to this activity. Notably, the contribution of these SLiMs differs between droplet dissolution and inhibition of droplet formation. For droplet dissolution, binding of PaRNase E to PaRhlB via the AR1 SLiM represents the minimal requirement, with RNA binding providing a secondary contribution (Fig. 5; compare AR1, AR4, and REER single mutants with wild type). Importantly, the RNA-binding affinity of the RNase E AR1 mutant is similar to that of the wild-type (37), supporting the necessity of RNase E–PaRhlB interaction for droplets dissolution. Consistent with this, *E. coli* RNase E binds RNA but does not interact with PaRhlB and fails to dissolve PaRhlB droplets, demonstrating that RNA binding alone is insufficient to drive this transition. Inhibition of droplet formation instead requires the presence of at least one of the three SLiMs (Fig. 6), indicating that the molecular requirements for RNase E^C-ter^ inhibition differ depending on whether droplets are already formed or are prevented from assembling. These differences may reflect changes in RNA or PaRhlB concentrations within droplets.

A distinct behaviour was observed for the *P. aeruginosa* RNA helicases RhlE1 and RhlE2. In contrast to PaRhlB, the formation and stability of RhlE1 and RhlE2 droplets were not influenced by the PaRNase E^C-ter^. Previously performed RNA-binding assays showed that RhlE1 and RhlE2 associate with RNA with substantially higher affinity than PaRhlB. Specifically, when RNA binding assays were performed using a 5′-overhang dsRNA_26/14_ substrate in the absence of ATP and identical experimental conditions, RhlE1 and RhlE2 exhibited dissociation constants of 2.7 nM and 1.5 nM, respectively, compared with 55 nM for PaRhlB ((26), and this work). This higher RNA affinity, together with the absence of interaction with the RNase E scaffolding region, likely contributes to the reduced susceptibility of RhlE1 and RhlE2 droplets to disruption relative to PaRhlB.

We previously found that RhlE2 can also interact with RNase E, but through an RNA-dependent mechanism involving the N-terminal catalytic domain of RNase E (47). This suggests that PaRhlB and RhlE2 might route their RNA substrates into different regulatory pathways: RhlB-dependent RNAs appear to be processed through the RNaseE^C-ter^-dependent (i.e. RNA degradosome-dependent) pathway, whereas RhlE2 RNA substrates may be handled by a separate, RNaseE^C-ter^-independent mechanism. We are currently testing this model by identifying the direct RNA targets of RhlB and RhlE2 and determining whether their regulation depends on RNase E interaction.

RNase E–mediated dissolution of PaRhlB condensates has physiological consequences during growth at low temperature. Low temperatures stabilise RNA secondary structures that can impede mRNA translation, RNA processing/turnover or ribosome biogenesis, thereby increasing the requirement for RNA helicase–mediated RNA remodelling (48). Consistent with this, RhlB contributes to efficient growth at low temperature in *Yersinia pseudotuberculosis* (49), whereas *E. coli* and *Pseudomonas syringae* Lz4W depend on other RNA helicases to support cold adaptation (43,50,51). Here, we show that in *P. aeruginosa* PaRhlB is required for optimal growth during cold adaptation, but also that its overexpression is detrimental. This deleterious effect on growth depends on the NTE of PaRhlB and not on its catalytic activity and most likely reflects misregulation of essential RNAs that could be sequestered in PaRhlB condensates; however, this remain to be verified. Importantly, RNase E^C-ter^ counteracts this effect, probably by preventing RhlB condensate persistence and, in general, by maintaining a correct RNA metabolism during cold growth. Type II RhlB homologs occur predominantly in environmental and marine Gammaproteobacteria (e.g., *Halopseudomonas*, *Spongiibacter*, *Marinobacter*, *Moraxella*, *Acinetobacter*), with additional representatives in sulfate-reducing *Desulfobacterota*/*Desulfovibrionia* lineages (e.g., *Desulfobacter latus*, *Desulfurhabdus amnigena*, *Desulforuvibrio alkaliphilus*). Functional validation in representative non-Pseudomonas species could determine the extent to which this regulatory mechanism is conserved and how it contributes to species-specific adaptation of RNA metabolism.

Beyond the RhlB-RNase E system, our findings confirm LLPS as an important layer of control in cellular RNA metabolism. Many bacterial and eukaryotic RNA helicases contain IDRs. In bacteria, for example, several *E. coli* helicases (Csda, DbpA, and RhlE) harbour IDR, undergo LLPS *in vitro*, and form foci *in vivo* when overexpressed (52,53). In the cyanobacterium *Synechocystis* sp. strain PCC 6803 the RNA helicase CrhR compartmentalizes into dynamic condensates in response to abiotic stress (24). In eukaryotes, among others, DDX3X, DDX10 and DDX6 provide illustrative examples within a much broader landscape. DDX3X uses disordered regions at both termini to enhance RNA unwinding and to recognize structured substrates such as G-quadruplexes (54–56). DDX10, in turn, requires its flanking IDRs to form nucleolar condensates and sustain pre-rRNA processing (57). DDX6 associates with mRNAs of low GC content, promotes their recruitment into processing bodies, and influences metabolic plasticity and chemoresistance (58). These proteins exemplify a larger class of eukaryotic RNA helicases whose IDRs confer substrate selectivity, spatial organization, and regulatory flexibility. Controlling RNA helicase condensate formation could therefore represent a widespread regulatory strategy for tuning RNA metabolism in diverse biological systems.

## Materials and Methods

### Bacteria growing conditions

*P. aeruginosa* PAO1 was routinely cultured at 37 °C in nutrient yeast broth (NYB; 25 g Difco nutrient broth and 5 g Difco yeast extract per liter) with agitation at 180 rpm, or on nutrient agar (NA; 40 g Oxoid blood agar base and 5 g Difco yeast extract per liter), unless stated otherwise. *E. coli* strains were grown under comparable conditions in LB broth or on LB agar. When appropriate, antibiotics were included at the following final concentrations: ampicillin (100 µg/mL), kanamycin (50 µg/mL), tetracycline (20 µg/mL), and gentamicin (10 µg/mL) for *E. coli*; and tetracycline (50 or 100 µg/mL) and gentamicin (50 µg/mL) for *P. aeruginosa*.

### Cloning procedures

The strains and plasmids used in this study are listed in Table S4. Plasmids were constructed either by restriction enzyme digestion followed by ligation with T4 DNA ligase, or by using the NEBuilder HiFi DNA Assembly Kit, according to the manufacturer’s instructions or standard protocols. The specific cloning procedure for each plasmid is detailed in Table S5. DNA fragments and vector backbones used for DNA assembly were amplified by PCR with primers listed in Table S6. Ligation mixtures or DNA assembly reactions were transformed into chemically competent *E. coli* DH5α cells by heat shock and plated on the appropriate selective LB agar medium. The resulting colonies were screened by PCR for the presence of the desired insert. Following plasmid extraction (Miniprep Kit, Sigma-Aldrich/Merck), plasmids were verified by Sanger sequencing (Microsynth).

To construct chromosomal *rhl* gene deletion or insert *mRuby* in frame with *rhl*, the suicide vectors pEXG2ΔrhlB or pME3087_RhlB-mRuby2 were used. The plasmids were introduced into *P. aeruginosa* PAO1 strain by triparental mating using the *E. coli* strain HB101 (pRK2013). After selection with the appropriate antibiotic (tetracycline for pME3087 or gentamicin for pEXG2), merodiploids were resolved as described in (37). Insertion/deletion were identified by colony PCR.

### Protein purification

His_10_Smt3-tagged PaRhlB, PaRhlB^K169A^, PaRhlB^90-507^, PaRhlB^111-507^, PaRhlB^1-111^, PaRhlB-msfGFP, RhlB^111-507^-msfGFP, PaRhlB^1-111^-msfGFP and *E. coli* RhlB (EcRhlB) proteins were purified from soluble bacterial lysates by nickel-agarose affinity chromatography as described previously in the supplementary information of Hausmann *et al.* 2021. The recombinant His_10_Smt3-tagged proteins were recovered predominantly in the 250 mM imidazole fraction. The imidazole elution profiles were monitored by SDS-PAGE (250 mM imidazole fraction for every recombinant protein is shown in Fig. S3). Imidazole was removed by overnight dialysis at 4°C in buffer A (50 mM Tris-HCl, pH 8.0, 500 mM NaCl, 10% glycerol). Otherwise stated the His_10_-Smt3 tag was not cleaved. For assays in Fig. S4, the His_10_-Smt3 tag was cleaved by the specific protease ULP-1 as described in the supplementary information of Hausmann *et al.* 2021. For LLPS experiments (see below), the 250 mM imidazole fraction was concentrated with a centrifugal filter (Amicon Ultra 4; 10’000 MWCO) and then stored at -80°C. Protein concentration was determined by using the Bio-Rad dye reagent with BSA as the standard.

*P. aeruginosa* RNase E^572-11057^ (named PaRNase E^C-ter^ or PaRNE^C-ter^) and *E. coli* RNase E^588-1061^ (named EcRNase E^C-ter^ or and EcRNE^C-ter^) carrying a His_10_Smt3 tag at the N-terminus and a Twin-Strep-Flag-tag at the C-terminus, were purified as described in (37). Briefly, RNase E derivatives were purified as described above. The 250 mM imidazole fraction was applied to a 1 ml of Strep-Tactin Sepharose (Iba) column that had been equilibrated with buffer C (50 mM Tris pH8, 200 mM NaCl, 10% glycerol). The column was washed with 5 ml of buffer C and then eluted stepwise with 1 ml aliquots of buffer C containing, 2.5 mM D-desthiobiotin respectively. The elution profiles were monitored by SDS-PAGE. All purified proteins used in biochemical assays throughout this study are shown in Fig. S3. Note that PaRNase E^C-ter^ mutated variants are purified as described above and are shown in the supplemental figure S6 of (37). The AR1mut, REERmut, and AR4mut alanine mutants are described in Table S4 of Geslain et al. (2025). Briefly, AR1mut replaces 12 Arg residues with Ala; AR4mut replaces 8 Arg residues and 2 Gln residues with Ala; and REERmut replaces 41 Arg residues with Ala.

### In vitro liquid–liquid phase separation (LLPS) assays

All LLPS-based assays were performed at 25 °C using proteins pre-diluted in buffer H (50 mM HEPES, pH 7.2–7.5; 500 mM NaCl; 0.01% Triton X-100). Unless otherwise stated, protein mixtures consisted of the indicated RhlB variant together with its msfGFP-tagged version at a 100:1 molar ratio (unlabeled:msfGFP-labeled). Droplet formation was quantified using Fiji (59) by measuring the fraction of the well area occupied by droplets or droplets fluorescence intensity as compared to the background. Below are details for each specific LLPS assay:

#### A. Standard LLPS assay

Reactions (25 µl total) were assembled in 1.5 mL Eppendorf tubes by adding 5 µL of diluted protein solution to 20 µl of reaction mix to yield final concentrations of: 50 mM HEPES (pH 7.2–7.5), 0.002% Triton X-100, 0.4 µg/µl BSA, 2 mM MgCl₂, 100 mM NaCl, and 250 ng/µl poly(U) (or RNA, as specified). Samples were incubated for 10 min at 25 °C, transferred to a 384-well plate (Starlab, 4TI-0214), and incubated for an additional 50 min before imaging with an EVOS FL microscope (Life Technologies) at 40× magnification. In experiments without msfGFP-tagged protein, imaging was performed using a Nikon Ts2R-FL microscope (40× magnification).

#### B. Effect of RNase E on droplet formation

For assays testing *P. aeruginosa* RNase E (PaRNase E) or *E. coli* RNase E (EcRNase E), the RNase E variant and RhlB were pre-diluted together in buffer H at a 1:1 molar ratio before assembly as described in standard LLPS assay above.

#### C. Droplet dissolution assay

Initial reaction mixtures (12.5 µl) containing 50 mM HEPES (pH 7.2–7.5), 0.002% Triton X-100, 0.4 µg/µl BSA, 2 mM MgCl_2_, 100 mM NaCl, and 250 ng/µl poly(U) RNA (or RNA as specified), and protein (as specified) were prepared by adding 2.5 µL of diluted protein to 10 µl of reaction mix. Final concentrations in this initial volume were: 50 mM HEPES (pH 7.2–7.5), 0.002% Triton X-100, 0.4 µg/µL BSA, 2 mM MgCl_2_, 100 mM NaCl, 250 ng/µL poly(U) (or RNA, as specified), and 1.25 µM RhlB plus its msfGFP-tagged version at a 100:1 molar ratio. Samples were incubated for 60 min at 25 °C. Subsequently, 12.5 µl of solution containing 50 mM HEPES (pH 7.2–7.5), 0.002% Triton X-100, 0.4 µg/µl BSA, 2 mM MgCl_2_, 100 mM NaCl, 250 ng/µl poly(U), 1.25 µM RhlB, and 2.5 µM RNase E as specified (or buffer H for controls) was added. After 10 min at 25 °C, samples were transferred to a 384-well plate and incubated for an additional 50 min before imaging with an EVOS FL microscope (Life Technologies) at 40× magnification.

### HDX-MS assays

HDX-MS experiments were performed at the UniGe Protein Biochemistry Platform (University of Geneva, Switzerland) following a well-established protocol with minimal modifications (60). Tables S2 and S3 detail all the reaction conditions and analytical values (raw data; H/D exchange levels and HDX differences worksheets). The following conditions were tested and compared: i) RhlB, and ii) RhlB + RNase E. Briefly, a first concentrated mix of 8 μl was prepared and preincubated for 5 min at 0 °C before initiating deuteration reaction. Deuterium exchange reaction was initiated by adding deuterated buffer at 0 °C to a final reaction volume of 50 μl. Reactions were carried out for 3 sec, 30 sec and 5 min and terminated by the sequential addition of ice-cold quench buffer. Samples were immediately frozen in liquid nitrogen and stored at -80 °C for up to two weeks. Number of replicates for each condition are described in Table S2 and raw data in Table S3.

To quantify deuterium uptake into the protein, samples were thawed and injected in a UPLC system immersed in ice with 0.1 % FA as liquid phase. The protein was digested via an immobilized Pepsin column (AffiPro AP-PC-001), and peptides were collected onto a Nucleodur 300-5 C18 metal-beads pre-column filter (Macherey-Nagel). The trap was subsequently eluted, and peptides separated with a C18, 175 Å, 1.9 μm particle size Thermo Hypersil Gold Vanquish 100 x 2.1 mm column over a gradient of 8 – 30 % buffer C over 20 min at 150 µl/min (Buffer B: 0.1% formic acid; buffer C: 100% acetonitrile). Mass spectra were acquired on an Orbitrap Velos Pro (Thermo), for ions from 400 to 2200 m/z using an electrospray ionization source operated at 300 °C, 5 kV of ion spray voltage. Peptides were identified by data-dependent acquisition of a non-deuterated sample after MS/ MS and data were analyzed by Mascot 2.6 using a database composed of purified proteins and known contaminants. Precursor mass tolerance was set to 10 ppm and fragment mass tolerance to 0.6 Da. Protein digestion was set as nonspecific. All peptides analysed are shown in tables S3a-c. Deuterium incorporation levels were quantified using HD examiner version 3.4.2 software (Sierra Analytics), and quality of every peptide was checked manually. Results are presented as percentage of maximal deuteration compared to theoretical maximal deuteration. Criteria to define a change in deuteration level between two states as significant is indicated in the table (change must be >10% and >1 Da. When triplicates are obtained, p value of student t-test must be < 0.01).

### ATPase activity assays

Reaction mixtures (15 µl) containing 50 mM HEPES (pH 7.2–7.5), 5 mM DTT, 2 mM MgGlu, 1 mM ATP, 50 mM KGlu, 1.5 U RNasin plus (Promega), RNA (none, 250 ng/µl poly(U), 125 ng/µl total RNA, 5 µM 5’overhang_dsRNA_26/14_, 5 µM ssRNA), and protein (as specified) were incubated for 15 minutes at 37 °C. Reactions were quenched by adding 3.8 µl of 5 M formic acid. Subsequently, 200 µl, 500 µl, or 1 ml of malachite green reagent (BIOMOL Green AK-111; Enzo) was added to the quenched reaction and thoroughly mixed. The mixtures were then incubated at room temperature for 30 minutes. A volume of 200 µl from each reaction was transferred to a 96-well transparent plate (TPP, 92096). Free phosphate was quantified by measuring absorbance at 620 nm (A₆₂₀) using a microplate reader (Spark, Tecan, Männedorf, Switzerland), and values were interpolated against a phosphate standard curve.

Alternatively, to assess the effect of buffer composition on intrinsic EcRhlB ATPase activity and its stimulation by RNase E, 15 µl reaction mixtures containing 50 mM Tris-HCl (pH 8.0), 1.5 U RNasin Plus (Promega), 100 mM KCl, 1 mM MgCl₂, 0.2 mM ATP, RNA (as specified above), and protein (as specified) were incubated for 15 minutes at 37 °C. Released inorganic phosphate (Pi) was quantified as described above.

### RNA unwinding assays

RNA unwinding was monitored using a fluorescence-based assay as described in Hausmann et al. (2024). The RNA substrate 5’overhang_dsRNA_26/14_ (5’ovh_dsRNA_26 /14_), purchased from IDT, comprises a 14-base pair duplex region and a 12-nucleotide 5′ single-stranded overhang with the following sequence: 5′-AACAAAACAAAAUAGCACCGUAAAGC-3′-(IBRQ) / 5′-(Cy5)-GCUUUACGGUGCUA. The RNA substrate 3’overhang_dsRNA_26/14_ (3’ovh_dsRNA_26 /14_) comprises a 14-base pair duplex region and a 12-nucleotide 3′ single-stranded overhang with the following sequence: 5’-GCUUUACGGUGCUA-3’-[Cy5]/ [IBRQ]-5’-UAGCACCGUAAAGCAAACAAAACAAA-3’). The blunt RNA duplex substrate (dsRNA _14/14_) comprises a 14-base pair duplex region with the following sequence: ([Cy5]-5’-GCUUUACGGUGCUA-3’ / 5’-UAGCACCGUAAAGC-3’-[IBRQ]). In this setup, RNA unwinding is detected as an increase in Cy5 fluorescence upon separation of the labeled strand from its quencher-labeled complement.

Assays were performed at 37 °C in 384-well plates using a Victor Nivo plate reader (PerkinElmer, Basel, Switzerland), with excitation at 640 nm and emission measured at 685 nm. Each 15 μl reaction contained: 50 mM HEPES (pH 7.5), 50 mM KGlu, 2 mM MgGlu, 1 mM ATP (or no ATP for control), 5 mM DTT, 50 nM RNA duplex (as specified), 3 μM RNA competitor (5′-GCUUUACGGUGCUA), 1.2 U/μl RNasin Plus (Promega), and purified recombinant protein as specified. The RNA competitor was included to prevent re-annealing of the Cy5-labeled strand to its quencher-labeled complement. Fluorescence measurements began one minute after protein addition and were recorded over time. Data were baseline-corrected by subtracting the fluorescence of a ‘no helicase’ control included in each experiment. To calculate the fraction of RNA unwound per minute, fluorescence values were normalized against the signals from fully dissociated (maximum fluorescence) and fully annealed (minimum fluorescence) RNA. The maximum fluorescence was determined from reactions containing 500 nM RhlE2, and this value corresponded closely with that obtained by heat-denaturing the RNA substrate at 90 °C for 5 minutes (Hausmann et al., 2024). Unwinding rate constants were estimated from the linear portion of the fluorescence-time curves (see Fig. 2B).

### RNA binding affinity assays

RNA binding was measured by fluorescence polarization (FP) assay as previously described in Hausmann et al., 2024. RNA was purchased from IDT (LubioScience GmbH, Zurich, Switzerland). Assays were performed in 384-well black plates (Corning 3820). Reaction mixtures (20 µl) containing 50 mM HEPES (pH 7.2–7.5), 5 mM DTT, 2 mM MgGlu, 1 mM of the non-hydrolysable ATP analogue adenosine-5′-(β,γ-imido)triphosphate (AMP-PNP), 50 mM KGlu, 1 nM 5′-FAM-labeled duplex RNA with a 5’ 12-nucleotide overhang (5′ovh_dsRNA26/14; 5′-FAM-AACAAAACAAAAUAGCACCGUAAAGC/5′-GCUUUACGGUGCUA), and protein (at concentrations as specified) were incubated at 25 °C for 15 minutes.

Fluorescence anisotropy measurements were performed using a Victor Nivo plate reader (PerkinElmer, Basel, Switzerland). The excitation wavelength was set to 480 nm, and emission was measured at 530 nm. Data points were collected in triplicate for each sample. The measured anisotropy values were normalized relative to the protein-free control. Data represent the mean ± standard deviation of six independent experiments.

To determine binding constants, changes in fluorescence polarization were plotted against protein concentration, and binding curves were fitted using the “One site-specific binding” model in GraphPad Prism (version 10.5, GraphPad Software) as described by the following equation:

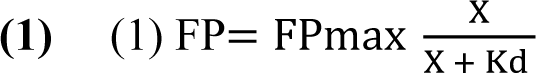

where X is the protein concentration, Kd the equilibrium dissociation constant.

### Bacterial two-hybrid assays

Bacterial two-hybrid experiments were performed as previously described (47). Briefly, recombinant pKT25 and pUT18C derivative plasmids were transformed simultaneously into the *E. coli* BTH101 strain. Transformants were spotted onto McConkey agar plates supplemented with 1 mM isopropyl β-d-thiogalactoside (IPTG) in the presence of 100 μg/ml ampicillin, 50 μg/ml kanamycin. Positive interactions were identified as red colonies after 24h incubation at 30°C. Three independent experiments were performed, and representative plates are shown.

## Acknowledgements

We would like to thank Sandra Amandine Marie Geslain for preliminary BTH experiments and helpful discussion. We also thank Rémy Visentin from the Proteins and Peptides platform (University of Geneva) for technical help with HDX-MS experiments.

## Notes

### Competing Interest Statement

The authors have declared no competing interest.

### Funding

- Swiss National Science Foundation: PCEFP3_203343

## References

1. Linder, P. and Jankowsky, E. (2011) From unwinding to clamping - the DEAD box RNA helicase family. Nat Rev Mol Cell Biol, 12, 505–516.

2. Jarmoskaite, I. and Russell, R. (2014) RNA helicase proteins as chaperones and remodelers. Annu Rev Biochem, 83, 697–725.

3. Hilbert, M., Karow, A.R. and Klostermeier, D. (2009) The mechanism of ATP-dependent RNA unwinding by DEAD box proteins. Biol Chem, 390, 1237–1250.

4. Caruthers, J.M. and McKay, D.B. (2002) Helicase structure and mechanism. Curr Opin Struct Biol, 12, 123–133.

5. Fairman-Williams, M.E., Guenther, U.P. and Jankowsky, E. (2010) SF1 and SF2 helicases: family matters. Curr Opin Struct Biol, 20, 313–324.

6. Theissen, B., Karow, A.R., Köhler, J., Gubaev, A. and Klostermeier, D. (2008) Cooperative binding of ATP and RNA induces a closed conformation in a DEAD box RNA helicase. Proc Natl Acad Sci U S A, 105, 548–553.

7. Chen, Y., Potratz, J.P., Tijerina, P., Del Campo, M., Lambowitz, A.M. and Russell, R. (2008) DEAD-box proteins can completely separate an RNA duplex using a single ATP. Proceedings of the National Academy of Sciences, 105, 20203–20208.

8. Hamann, F., Enders, M. and Ficner, R. (2019) Structural basis for RNA translocation by DEAH-box ATPases. Nucleic Acids Res, 47, 4349–4362.

9. Singleton, M.R., Dillingham, M.S. and Wigley, D.B. (2007) Structure and mechanism of helicases and nucleic acid translocases. Annu Rev Biochem, 76, 23–50.

10. Gilman, B., Tijerina, P. and Russell, R. (2017) Distinct RNA-unwinding mechanisms of DEAD-box and DEAH-box RNA helicase proteins in remodeling structured RNAs and RNPs. Biochem Soc Trans, 45, 1313–1321.

11. Donsbach, P. and Klostermeier, D. (2021) Regulation of RNA helicase activity: principles and examples. Biol Chem, 402, 529–559.

12. Sloan, K.E. and Bohnsack, M.T. (2018) Unravelling the Mechanisms of RNA Helicase Regulation. Trends Biochem Sci, 43, 237–250.

13. Choi, J.M., Holehouse, A.S. and Pappu, R.V. (2020) Physical Principles Underlying the Complex Biology of Intracellular Phase Transitions. Annu Rev Biophys, 49, 107–133.

14. Borcherds, W., Bremer, A., Borgia, M.B. and Mittag, T. (2021) How do intrinsically disordered protein regions encode a driving force for liquid-liquid phase separation? Curr Opin Struct Biol, 67, 41–50.

15. Nandana, V. and Schrader, J.M. (2021) Roles of liquid-liquid phase separation in bacterial RNA metabolism. Curr Opin Microbiol, 61, 91–98.

16. Dai, Z. and Yang, X. (2024) The regulation of liquid-liquid phase separated condensates containing nucleic acids. FEBS J, 291, 2320–2331.

17. Hyman, A.A., Weber, C.A. and Julicher, F. (2014) Liquid-liquid phase separation in biology. Annu Rev Cell Dev Biol, 30, 39–58.

18. Hansma, H.G. (2017) Better than Membranes at the Origin of Life? Life (Basel), 7.

19. Hooper, C. and Hilliker, A. (2013) Packing them up and dusting them off: RNA helicases and mRNA storage. Biochim Biophys Acta, 1829, 824–834.

20. Weis, K. (2021) Dead or alive: DEAD-box ATPases as regulators of ribonucleoprotein complex condensation. Biol Chem, 402, 653–661.

21. Overwijn, D. and Hondele, M. (2023) DEAD-box ATPases as regulators of biomolecular condensates and membrane-less organelles. Trends Biochem Sci, 48, 244–258.

22. Dörner, K. and Hondele, M. (2024) The Story of RNA Unfolded: The Molecular Function of DEAD- and DExH-Box ATPases and Their Complex Relationship with Membraneless Organelles. Annu Rev Biochem, 93, 79–108.

23. Khemici, V. and Linder, P. (2016) RNA helicases in bacteria. Curr Opin Microbiol, 30, 58–66.

24. Whitman, B.T., Wang, Y., Murray, C.R.A., Glover, M.J.N. and Owttrim, G.W. (2023) Liquid-Liquid Phase Separation of the DEAD-Box Cyanobacterial RNA Helicase Redox (CrhR) into Dynamic Membraneless Organelles in Synechocystis sp. Strain PCC 6803. Appl Environ Microbiol, 89, e0001523.

25. Fuller-Pace, F.V., Nicol, S.M., Reid, A.D. and Lane, D.P. (1993) DbpA: a DEAD box protein specifically activated by 23s rRNA. Embo j, 12, 3619–3626.

26. Hausmann, S., Geiser, J., Allen, G.E., Geslain, S.A.M. and Valentini, M. (2024) Intrinsically disordered regions regulate RhlE RNA helicase functions in bacteria. Nucleic Acids Res, 52, 7809–7824.

27. Py, B., Higgins, C.F., Krisch, H.M. and Carpousis, A.J. (1996) A DEAD-box RNA helicase in the *Escherichia coli* RNA degradosome. Nature, 381, 169–172.

28. Carpousis, A.J. (2007) The RNA degradosome of *Escherichia coli*: an mRNA-degrading machine assembled on RNase E. Annu Rev Microbiol, 61, 71–87.

29. Vanzo, N.F., Li, Y.S., Py, B., Blum, E., Higgins, C.F., Raynal, L.C., Krisch, H.M. and Carpousis, A.J. (1998) Ribonuclease E organizes the protein interactions in the *Escherichia coli* RNA degradosome. Genes Dev, 12, 2770–2781.

30. Coburn, G.A., Miao, X., Briant, D.J. and Mackie, G.A. (1999) Reconstitution of a minimal RNA degradosome demonstrates functional coordination between a 3’ exonuclease and a DEAD-box RNA helicase. Genes Dev, 13, 2594–2603.

31. Khemici, V., Toesca, I., Poljak, L., Vanzo, N.F. and Carpousis, A.J. (2004) The RNase E of *Escherichia coli* has at least two binding sites for DEAD-box RNA helicases: functional replacement of RhlB by RhlE. Mol Microbiol, 54, 1422–1430.

32. Liou, G.G., Chang, H.Y., Lin, C.S. and Lin-Chao, S. (2002) DEAD box RhlB RNA helicase physically associates with exoribonuclease PNPase to degrade double-stranded RNA independent of the degradosome-assembling region of RNase E. J Biol Chem, 277, 41157–41162.

33. Bruce, H.A., Du, D., Matak-Vinkovic, D., Bandyra, K.J., Broadhurst, R.W., Martin, E., Sobott, F., Shkumatov, A.V. and Luisi, B.F. Analysis of the natively unstructured RNA/protein-recognition core in the *Escherichia coli* RNA degradosome and its interactions with regulatory RNA/Hfq complexes.

34. Chandran, V., Poljak, L., Vanzo, N.F., Leroy, A., Miguel, R.N., Fernandez-Recio, J., Parkinson, J., Burns, C., Carpousis, A.J. and Luisi, B.F. (2007) Recognition and cooperation between the ATP-dependent RNA helicase RhlB and ribonuclease RNase E. Journal of molecular biology, 367, 113–132.

35. Worrall, J.A., Howe, F.S., McKay, A.R., Robinson, C.V. and Luisi, B.F. (2008) Allosteric activation of the ATPase activity of the *Escherichia coli* RhlB RNA helicase. J Biol Chem, 283, 5567–5576.

36. Khemici, V., Poljak, L., Toesca, I. and Carpousis, A.J. (2005) Evidence in vivo that the DEAD-box RNA helicase RhlB facilitates the degradation of ribosome-free mRNA by RNase E. Proc Natl Acad Sci U S A, 102, 6913–6918.

37. Geslain, S.A.M., Hausmann, S., Geiser, J., Allen, G.E., Gonzalez, D. and Valentini, M. (2025) Critical functions and key interactions mediated by the RNase E scaffolding domain in Pseudomonas aeruginosa. PLoS genetics, 21, e1011618.

38. Aït-Bara, S., Carpousis, A.J. and Quentin, Y. (2015) RNase E in the γ-Proteobacteria: conservation of intrinsically disordered noncatalytic region and molecular evolution of microdomains. Mol Genet Genomics, 290, 847–862.

39. Tejada-Arranz, A., de Crécy-Lagard, V. and de Reuse, H. (2020) Bacterial RNA Degradosomes: Molecular Machines under Tight Control. Trends Biochem Sci, 45, 42–57.

40. Kumari, H., Murugapiran, S.K., Balasubramanian, D., Schneper, L., Merighi, M., Sarracino, D., Lory, S. and Mathee, K. (2014) LTQ-XL mass spectrometry proteome analysis expands the Pseudomonas aeruginosa AmpR regulon to include cyclic di-GMP phosphodiesterases and phosphoproteins, and identifies novel open reading frames. J Proteomics, 96, 328–342.

41. Hardwick, S.W., Chan, V.S., Broadhurst, R.W. and Luisi, B.F. (2011) An RNA degradosome assembly in *Caulobacter crescentus*. Nucleic Acids Res, 39, 1449–1459.

42. Uversky, V.N., Gillespie, J.R. and Fink, A.L. (2000) Why are “natively unfolded” proteins unstructured under physiologic conditions? Proteins, 41, 415–427.

43. Iost, I. and Dreyfus, M. (2006) DEAD-box RNA helicases in Escherichia coli. Nucleic Acids Res, 34, 4189–4197.

44. Huen, J., Lin, C.-L., Golzarroshan, B., Yi, W.-L., Yang, W.-Z. and Yuan, H.S. (2017) Structural Insights into a Unique Dimeric DEAD-Box Helicase CshA that Promotes RNA Decay. Structure, 25, 469–481.

45. Zetzsche, H., Raschke, L. and Furtig, B. (2023) Allosteric activation of RhlB by RNase E induces partial duplex opening in substrate RNA. Front Mol Biosci, 10, 1139919.

46. Al-Husini, N., Tomares, D.T., Bitar, O., Childers, W.S. and Schrader, J.M. (2018) α-Proteobacterial RNA Degradosomes Assemble Liquid-Liquid Phase-Separated RNP Bodies. Molecular cell, 71, 1027–1039.e1014.

47. Hausmann, S., Gonzalez, D., Geiser, J. and Valentini, M. (2021) The DEAD-box RNA helicase RhlE2 is a global regulator of Pseudomonas aeruginosa lifestyle and pathogenesis. Nucleic Acids Res, 49, 6925–6940.

48. Phadtare, S. and Severinov, K. (2010) RNA remodeling and gene regulation by cold shock proteins. RNA Biology, 7, 788–795.

49. Jiang, X., Keto-Timonen, R., Skurnik, M. and Korkeala, H. (2019) Role of DEAD-box RNA helicase genes in the growth of Yersinia pseudotuberculosis IP32953 under cold, pH, osmotic, ethanol and oxidative stresses. PLoS One, 14, e0219422.

50. Hussain, A. and Ray, M.K. (2024) Role of DEAD-box RNA helicases in low-temperature adapted growth of Antarctic Pseudomonas syringae Lz4W. Microbiol Spectr, 12, e0433522.

51. Awano, N., Xu, C., Ke, H., Inoue, K., Inouye, M. and Phadtare, S. (2007) Complementation Analysis of the Cold-Sensitive Phenotype of the *Escherichia coli csdA* Deletion Strain. Journal of Bacteriology, 189, 5808–5815.

52. Hondele, M., Sachdev, R., Heinrich, S., Wang, J., Vallotton, P., Fontoura, B.M.A. and Weis, K. (2019) DEAD-box ATPases are global regulators of phase-separated organelles. Nature, 573, 144–148.

53. Weis, K. and Hondele, M. (2022) The Role of DEAD-Box ATPases in Gene Expression and the Regulation of RNA-Protein Condensates. Annu Rev Biochem, 91, 197–219.

54. Floor, S.N., Condon, K.J., Sharma, D., Jankowsky, E. and Doudna, J.A. (2016) Autoinhibitory Interdomain Interactions and Subfamily-specific Extensions Redefine the Catalytic Core of the Human DEAD-box Protein DDX3. J Biol Chem, 291, 2412–2421.

55. He, Y.N., Han, X.R., Wang, D., Hou, J.L. and Hou, X.M. (2024) Dual mode of DDX3X as an ATP-dependent RNA helicase and ATP-independent nucleic acid chaperone. Biochem Biophys Res Commun, 714, 149964.

56. Toyama, Y., Takeuchi, K. and Shimada, I. (2025) Regulatory role of the N-terminal intrinsically disordered region of the DEAD-box RNA helicase DDX3X in selective RNA recognition. Nat Commun, 16, 7762.

57. Wang, X., Hu, G., Wang, L., Lu, Y., Liu, Y., Yang, S., Liao, J., Zhao, Q., Huang, Q., Wang, W. et al. (2024) DEAD-box RNA helicase 10 is required for 18S rRNA maturation by controlling the release of U3 snoRNA from pre-rRNA in embryonic stem cells. Nat Commun, 15, 10303.

58. Bi, H., Li, W., Ren, L., Zhang, H., Dong, L., Chan, A., Wang, X., Yang, L., Zhang, X., Xue, M. et al. (2025) DDX6 undergoes phase separation to modulate metabolic plasticity and chemoresistance. Nat Commun.

59. Schindelin, J., Arganda-Carreras, I., Frise, E., Kaynig, V., Longair, M., Pietzsch, T., Preibisch, S., Rueden, C., Saalfeld, S., Schmid, B. et al. (2012) Fiji: an open-source platform for biological-image analysis. Nat Methods, 9, 676–682.

60. Wang, H., Lo, W.T., Vujičić Žagar, A., Gulluni, F., Lehmann, M., Scapozza, L., Haucke, V. and Vadas, O. (2018) Autoregulation of Class II Alpha PI3K Activity by Its Lipid-Binding PX-C2 Domain Module. Mol Cell, 71, 343–351.e344.

61. Jumper, J., Evans, R., Pritzel, A., Green, T., Figurnov, M., Ronneberger, O., Tunyasuvunakool, K., Bates, R., Žídek, A., Potapenko, A., et al. (2021) Highly accurate protein structure prediction with AlphaFold. Nature, 596, 583–589.

62. Schrodinger, LLC. (2015) The PyMOL Molecular Graphics System, Version 1.8.

63. García-Gutiérrez, C., Aparicio, T., Torres-Sánchez, L., Martínez-García, E., de Lorenzo, V., Villar, C.J. and Lombó, F. (2020) Multifunctional SEVA shuttle vectors for actinomycetes and Gram-negative bacteria. MicrobiologyOpen, 9, e1024.

