## Supplementary material for "RNase E resolves toxic condensates by counteracting phase separation of Type II RhlB helicases": SI

#### Supplementary Tables

**Table S1: List of unique peptides identified in the elution by LC/MS-MS.**

| Unique Peptide Sequence | Peptide start index | Peptide stop index | Domain |
|---|---|---|---|
| IFGKADGTQPQSGSSPVPQGPAGATAK | 24 | 50 | NTE |
| ADGTQPQSGSSPVPQGPAGATAK | 28 | 50 | NTE |
| GTAPTGEKPTEKRPR | 51 | 65 | NTE |
| KPRPPRADNESGAPR | 66 | 80 | NTE |
| ADNESGAPR | 72 | 80 | NTE |
| ERKPKPVDNWK | 112 | 122 | RecA1 |
| KPKPVDNWKLEDFVVEPQEGK | 114 | 134 | RecA1 |
| LEDFVVEPQEGK | 123 | 134 | RecA1 |
| LEDFVVEPQEGKTR | 123 | 136 | RecA1 |
| TAAFLISIITQLLQTPPPK | 186 | 204 | RecA1 |
| TAAFLISIITQLLQTPPPKER | 186 | 206 | RecA1 |
| ALIIAPTR | 213 | 220 | RecA1 |
| ELVVQIAK | 221 | 228 | RecA1 |
| ELVVQIAKDAAALTK | 221 | 235 | RecA1 |
| DAAALTKYTGLNVMTFVGGMDFDK | 229 | 252 | RecA1 |
| YTGLNVMTFVGGMDFDK | 236 | 252 | RecA1 |
| YTGLNVMTFVGGMDFDKQLK | 236 | 255 | RecA1 |
| FCDILVATPGR | 261 | 271 | RecA1 |
| GEVHLDMVEVMVLDEADR | 279 | 296 | RecA1 |
| GEVHLDMVEVMVLDEADRMLDMGFIPQVR | 279 | 307 | RecA1 |
| MLDMGFIPQVR | 297 | 307 | RecA1 |
| QTLLFSATFTDDVMNLAK | 320 | 337 | RecA1 |
| QWTVDPAIVEIEPENVASDTVEQHVYAVAGSDK | 338 | 370 | RecA2 |
| QWTVDPAIVEIEPENVASDTVEQHVYAVAGSDKYK | 338 | 372 | RecA2 |
| YKLLYNLVAQNNWER | 371 | 385 | RecA2 |
| LLYNLVAQNNWER | 373 | 385 | RecA2 |
| VMVFANRK | 386 | 393 | RecA2 |
| LTKDGISAAQMSGDVPQHK | 403 | 421 | RecA2 |
| DGISAAQMSGDVPQHK | 406 | 421 | RecA2 |
| DGISAAQMSGDVPQHKR | 406 | 422 | RecA2 |
| TLEGFREGK | 425 | 433 | RecA2 |
| IRVLVATDVAGR | 434 | 445 | RecA2 |
| VLVATDVAGR | 436 | 445 | RecA2 |
| GIHIDGISHVINFTLPEDPDDYVHR | 446 | 470 | RecA2 |
| AGASGTSISFAGEDDAFALPPIEELLGRK | 477 | 505 | RecA2 |
| KITCEMPPAELLKPVPR | 505 | 521 | RecA2 |
| ITCEMPPAELLKPVPR | 506 | 521 | RecA2 |

**Table S2: HDX experimental details**

| Description | PaRhlB | PaRhlB + PaRNase $E^{C\text{-}ter}$ |
|---|---|---|
| **Control sample** | Non-deuterated (ND) PaRhlB | Non-deuterated (ND) PaRhlB |
| **D2O incubation times (sec)** | 3s, 30s, 300s | 3s, 30s, 300s |
| **Replicates** | 3, 3, 2 | 3, 3, 3 |
| **Sequence coverage** | 75.0% | 75.0% |
| **Number of peptides analysed** | 149 | 149 |
| **Average peptide length** | 13 | 13 |
| **Average peptide redundancy** | 3.5 | 3.5 |
| **% D2O in Rxn** | 79 % | 79 % |
| **Temperature** | 0 °C | 0 °C |
| **Protein target (pmol)** | 615 | 603 |
| **Target concentration (µM)** | 12.3 | 12.1 |
| **Ligand concentration (µM)** | - | - |
| **Reaction volume** | 50 ul | 50 ul |
| **Protein mix volume (µl)** | 8 ul | 8 ul |
| **Quench buff. Vol (µl)** | 20 ul | 20 ul |
| **Deut buffer** | 20 mM Tris pH 8, 200 mM NaCl | |
| **Quench buffer** | 7 M UREA, 1 M NaCl, 100 mM NaH2PO4 (pH 2.4), 1 % Formic Acid (FA) | |
| **Criteria for HDX rate difference** | Difference of HDX level at a given timepoint is > 10 % and > 1 Da and p values of student t-test is < 0.01 | |

**Table S3: HDX raw data. See Excel file**

**Table S4: List of strains and plasmids**

| Strains | Genotype/relevant characteristics | Source |
|---|---|---|
| ***E.coli*** | | |
| Rosetta (DE3) | F- ompT hsdSB(rB- mB-) gal dcm (DE3) pRARE (CmR) | Novagen |
| DH5α | recA1 endA1 hsdR17 supE44 thi-1 gyrA96 relA1 Δ(lacZYA-argF)U169 [Φ80dlacZM15]F- NaIr | (1) |
| BTH101 | F-, cya-99, araD139, galE15, galK16, rpsL1 (Strr), hsdR2, mcrA1, mcrB1 | Euromedex |
| ***P. aeruginosa*** | | |
| PAO1 | Wild-type | (2) |
| PAE200 | PA01 containing a deletion of the *rhl* (PA3861) locus | this study |
| PAE29 | PA01 containing an insertion of mRuby2 with a flexible linker immediately upstream of the stop codon in *rhl* locus | this study |
| **Plasmids** | | |
| pET28b-10xHis-Smt3 | Broad host range vector for expression of N-terminal 10xHis-Smt3-tag proteins, KmR | (3) |
| pET28b-10xHis-Smt3-SF | Broad host range vector for expression of N-terminal 10xHis-Smt3-tag in N-terminal and Strep-Flag-tag in C-terminal proteins, KmR | (3) |
| pKT25 | Two-hybrid plasmid, cyaAT25 fusion, KmR | (4) |
| pUT18C | Two-hybrid plasmid, cyaAT18 fusion, ApR | (4) |
| pME3087 | Suicide vector for allelic replacement; TcR; ColE1 replicon | (5) |
| pEXG2 | Suicide vector for allelic replacement; GmR; ColE1 ori, sacB | (6) |
| pSEVA4313 | Constitutive expression vector, SmR | (6) |
| pSEVA4313-Pbad | AraC-Pbad expression vector, SmR | (6) |
| pME6032 | lacIQ Ptac expression vector; GmR | (7) |
| pRK2013 | Helper plasmid; Tra+ KmR | (1) |
| pSmt3-RNAseE$^{572\text{-}1057}$-SF | pET28b-10xHis-Smt3 derivative for His-Smt3-RNAseE$^{572\text{-}1057}$-StrepFlag, KmR | (8) |
| pSmt3_RNase E$^{572\text{-}1057}$_AR4mut-SF | pET28b-10xHis-Smt3 derivative for His-Smt3-RNase E$^{572\text{-}1057}$_AR4mut-StrepFlag, KmR | (8) |
| pSmt3_ RNase E$^{572\text{-}1057}$_AR1mut-SF | pET28b-10xHis-Smt3 derivative for His-Smt3-RNase E$^{572\text{-}1057}$_AR1mut-StrepFlag, KmR | (8) |
| pSmt3_ RNase E$^{572\text{-}1057}$_REERmut-SF | pET28b-10xHis-Smt3 derivative for His-Smt3-RNase E$^{572\text{-}1057}$_REERmut-StrepFlag, KmR | (8) |
| pSmt3_ RNase E$^{572\text{-}1057}$_AR1mut AR4mut-SF | pET28b-10xHis-Smt3 derivative for His-Smt3- RNase E$^{572\text{-}1057}$_AR1mut AR4mut-StrepFlag, KmR | (8) |
| pSmt3_ RNase E$^{572\text{-}1057}$_AR1mut AR4mut REERmut-SF | pET28b-10xHis-Smt3 derivative for His-Smt3- RNase E$^{572\text{-}1057}$_AR1mut AR4mut REERmut-StrepFlag, KmR | (8) |
| pSmt3-RhlB | pET28b-10xHis-Smt3 derivative for His-Smt3-RhlB, KmR | (8) |
| pSmt3- RNase E$^{588\text{-}1061}$_SF | pET28b-10xHis-Smt3 derivative for His-Smt3-RNase E$^{588\text{-}1061}$-StrepFlag *E. coli*, KmR | This study |

| | | |
|---|---|---|
| pSmt3-RhlB$^{111-507}$ | pET28b-10xHis-Smt3 derivative for His-Smt3-RhlB$^{111-507}$, KmR | This study |
| pSmt3 RhlB$^{K169A}$ | pET28b-10xHis-Smt3 derivative for His-Smt3-RhlB$^{K169A}$, KmR | This study |
| pSmt3 RhlB-msfGFP | pET28b-10xHis-Smt3 derivative for His-Smt3-RhlB_msfGFP, KmR | This study |
| pSmt3 RhlB$^{111-507}$_msfGFP | pET28b-10xHis-Smt3 derivative for His-Smt3-RhlB$^{111-507}$_msfGFP, KmR | This study |
| pSmt3-RhlB$^{1-111}$ | pET28b-10xHis-Smt3 derivative for His-Smt3-RhlB RhlB$^{1-111}$, KmR | This study |
| pSmt3-RhlB$^{1-111}$_msfGFP | pET28b-10xHis-Smt3 derivative for His-Smt3-RhlB$^{1-111}$_msfGFP, KmR | This study |
| pSmt3-RhlB_Ec | pET28b-10xHis-Smt3 derivative for His-Smt3-RhlB*E. coli*, KmR | This study |
| pKT25-RhlB_RECA1 | pKT25 derivative for T25-RhlB_RECA1 expression, KmR | This study |
| pKT25-RhlB_RECA2 | pKT25 derivative for T25-RhlB_RECA2 expression, KmR | This study |
| pKT25-RhlB_RECA1+RECA2 | pKT25 derivative for T25-RhlB_RECA1+RECA2(111-507) expression, KmR | This study |
| pKT25-RhlB$^{1-111}$ | pKT25 derivative for T25-RhlB_NTE expression, KmR | This study |
| pKT25-RhlB | pKT25 derivative for T25-RhlB expression, KmR | (8) |
| pKT25-EcRhlB | pKT25 derivative for T25-RhlB of *E. coli* expression, KmR | (8) |
| pUT18C-RNAseE$^{733-793}$ | pUT18C derivative for T18-RNaseE$^{733-793}$ expression, ApR | (8) |
| pUT18C-RNAseE | pUT18C derivative for T18-RNaseE expression, ApR | (8) |
| pUT18C-RNAseE_Ec | pUT18C derivative for T18-RNaseE *E. coli* expression, ApR | This study |
| pSEVA4313-3F-RhlB | pSEVA4313 derivative for 3F-RhlB expression, Smr | This study |
| pSEVA4313-3F-RhlB$^{K169A}$ | pSEVA4313 derivative for 3F-RhlB$^{K169A}$ expression, Smr | This study |
| pSEVA4313-3F-RhlB$^{111-507}$ | pSEVA4313 derivative for 3F-RhlB$^{111-507}$ expression, Smr | This study |
| pME6032-3F-RhlB | pME6032 derivative for 3F-RhlB expression, GmR | This study |
| pSEVA4313-pBAD-RNAseE$^{572-1057}$_STREP | pSEVA4313-pBAD derivative for RNAseE$^{572-1057}$_STREP expression, SmR | This study |
| pEXG2Δ*rhlB* | pME3087 derivative for the deletion of the *rhl* gene, GmR | This study |
| pME3087-RhlB_mRuby2 | pME3087 derivative for the in-frame insertion of a mRuby2 sequence to the end of *rhl* gene, TcR | This study |

**Table S5: Cloning details**

| Name | Used for… | Backbone: | Insert(s) amplified with: | Template used in PCR | Type of cloning |
|---|---|---|---|---|---|
| pSmt3-EcRNAseE$^{588\text{-}1061}$_SF | Protein purification | pET28b-10xHis-Smt3-SF | p930-p931 | N3433 gDNA | Insert digested with BglII and NotI, vector digested with BamHI and NotI, ligated |
| pSmt3-RhlB$^{111\text{-}507}$ | Protein purification | pET28b-10xHis-Smt3 | p571-p499 | PA01 gDNA | Insert and vector digested with HindIII and XhoI, ligated |
| pSmt3-RhlB$^{K169A}$ | Protein purification | pET28b-10xHis-Smt3 | p629-p630 | pSmt3-RhlB | SDM |
| pSmt3-RhlB_msfGFP | Protein purification | pET28b-10xHis-Smt3 | p605-p498 | PA01gDNA | Insert and vector digested with HindIII and XhoI, ligated |
| pSmt3 RhlB$^{111\text{-}507}$-msfGFP | Protein purification | pET28b-10xHis-Smt3 | p605-p571 | PA01gDNA | Insert and vector digested with HindIII and XhoI, ligated |
| pSmt3-RhlB$^{1\text{-}111}$ | Protein purification | pET28b-10xHis-Smt3 | p690-p498 | PA01gDNA | Insert and vector digested with HindIII and XhoI, ligated |
| pSmt3-RhlB$^{1\text{-}111}$-msfGFP | Protein purification | pET28b-10xHis-Smt3-msfGFP | p688-p689 insert; p686-p687 vector | pSmt3_msfGFP; PA01gDNA | Gibson assembly |
| pSmt3-EcRhlB | Protein purification | pET28b-10xHis-Smt3 | p928-p929 | N3433gDNA | Insert and vector digested with HindIII and XhoI, ligated |
| pKT25-PaRhlB_REC1 | BTH assay | pKT25 | p811-p812 | PA01gDNA | Insert and vector digested with BamHI and EcoRI, ligated |
| pKT25-PaRhlB_REC2 | BTH assay | pKT25 | p813-p814 | PA01gDNA | Insert and vector digested with BamHI and EcoRI, ligated |
| pKT25-PaRhlB_REC1+REC2 | BTH assay | pKT25 | p811-p814 | PA01gDNA | Insert and vector digested with BamHI and EcoRI, ligated |
| pKT25-PaRhlB$^{1\text{-}111}$ | BTH assay | pKT25 | p889-p201 | PA01gDNA | Insert and vector digested with BamHI and EcoRI, ligated |
| pSEVA4313-3F-PaRhlB | Overexpression assay | pSEVA4313 | p799-p800 | pME6182-3F-RhlB | Insert and vector digested with EcoRI and HindIII, ligated |
| pSEVA4313-3F-PaRhlB$^{K169A}$ | Overexpression assay | pSEVA4313 | p799-p800 | pME6182-3F-RhlB_K169A | Insert and vector digested with EcoRI and HindIII, ligated |
| pSEVA4313-3F-PaRhlB$^{111\text{-}507}$ | Overexpression assay | pSEVA4313-3F | p911-p912 insert; p823-p824 vector | PA01gDNA; pSEVA4313-3F | Gibson assembly |
| pME6032-3F-RhlB | Inducible expression assay | pME6032 | p799-p499 insert | pSmt3-RhlB | Insert and vector digested with XhoI and EcoRI, ligated |
| pSEVA4313-pBAD-PaRNAseE$^{572\text{-}1057}$_STREP | Inducible expression assay | pSEVA4313-pBAD | p1001-p780 | pME6032-RNE-STREP | Insert and vector digested with SpeI and EcoRI, ligated |

| | | | | | |
|---|---|---|---|---|---|
| pEXG2ΔRhlB | Construction of PA01 strain | pEXG2 | p851-p852 insert; p537-p538 vector | PA01gDNA; pEXG2 | Gibson assembly |
| pME3087-RhlB_mRuby2 | Construction of PA01 strain | pME3087 | p372-p373; p374-p375; p376-p377 | PA01gDNA; pSw002-Pc-mRuby2 | Gibson assembly |

**Table S6: List pf primers**

| Primer | Primer sequence (5' -> 3') |
|---|---|
| p201 | GTACGGATCCCCTCAAAGCACTCAAGAAAATCTTCG |
| p372 | TAGGAGCATAGAATTCGAGGTGCACCTGGACATGG |
| p373 | GTTCTTCACCTTTGGAACCACCACCGTGCTTGCGCGGTACCG |
| p374 | CCGCGCAAGCACGGTGGTGGTTCCAAAGGTGAAGAACTGATC |
| p375 | CCATGCCTGCGTTCATTTATACAACTCATCCATACCA |
| p376 | GAGTTGTATAAATGAACGCAGGCATGGAGAAAAAAG |
| p377 | GAATGGCAAAAGCTTGCATCGCAGAGCAGCAAGG |
| p483 | GGTGGTGGTTCCCGTAA |
| p498 | CAGTAAGCTTCAGTGCTCAAAGCACTCAAGAAAATC |
| p499 | CATGCTCGAGTCAGTGCTTGCGCGGTAC |
| p537 | GGATCCTCTAGAGTCGACCTGC |
| p538 | GAATTCGGTACCTTAATTA |
| p571 | CAGTAAGCTTCAGTGGTCGAACCGCAGGAAG |
| p605 | CATGCTCGAGTTATTTGTAGAGTTCATCCATGCCGT |
| p629 | CAGACCGGCACCGGCGCGACCGCCGCGTTCCTC |
| p630 | GAGGAACGCGGCGGTCGCGCCGGTGCCGGTCTG |
| p686 | TGAAGCTTGTCGACGGAGCT |
| p687 | GGTGGTGGTTCCCGTAAAGG |
| p688 | TCCGTCGACAAGCTTCAGTGCTCAAAGCACTCAAGAAAATCT |
| p689 | TACGGGAACCACCACCCACGAAGTCTTCCAGCTTCCAG |
| p690 | CATGCTCGAGTCACACGAAGTCTTCCAGCTTCCAG |
| p747 | GTCGACTCTAGAGGATCCGGTGACGCGGGCGA |
| p748 | TCGACCACGGCGATCTCGTAGTGGTTAAGGAA |
| p749 | GATCGCCGTGGTCGAACCGCAGGAAG |
| p750 | GAAATTAATTAAGGTACCGAATTCGGTCGGCTTCGTCGAGG |
| p780 | GATCACTAGTGTACCCGGGAGCTCTCATTTCTCGAAC |
| p797 | GTCGACTCTAGAGGATCCGTGCTCAAAGCACTCAAGAAAATCTTCG |
| p798 | GAAATTAATTAAGGTACCGAATTCCGACGTTCTCCGGCTCG |
| p799 | ACTGGAATTCAGAAGGAGATATACCATGGACTACAAAGACCATGACG |
| p800 | CAGTAAGCTTTCAGTGCTTGCGCGGTACC |
| p811 | ACGTGGATCCCGTGGTCGAACCGCAGGAAGG |
| p812 | ACTGGAATTCTTACTCCGGCTCGATCTCGACGAT |
| p813 | ACGTGGATCCCGTCGAGATCGAGCCGGAG |
| p814 | ACTGGAATTCTCAGTGCTTGCGCGGTAC |
| p823 | AAGCTTGCGGCCGCGTCGTG |
| p824 | CTTGTCATCGTCATCCTTGTAATCGATATCATGA |
| p851 | AGGTCGACTCTAGAGGATCCCAGTGAATTCCAGCTCGGCA |
| p852 | GAAATTAATTAAGGTACCGAATTCGTGCAAGCTTCGCAGAGC |
| p889 | GATCGAATTCTCACACGAAGTCTTCCAGCTTC |
| p928 | CAGTAAGCTTCAAGCAAAACACATTTAACAGAACAGAAG |
| p929 | GACTCTCGAGTTAACCTGAACGACGACGATTACG |
| p930 | ACTGAGATCTACCAAACCGACCGAGCAA |
| p931 | GTCAGCGGCCGCCTCAACAGGTTGCGGACG |
| p998 | TTAATTAAGGTACCGAATTCCGAGGAGGGGACTCGAACCC |
| p999 | AGGTCGACTCTAGAGGATCCGTGATCGACGCCTACCACCA |

| | |
|---|---|
| p1000 | CACAGGAAACAGAATTCATGGTGGTCGAACCGCAGG |
| p1001 | ACTGGAATTCAGAAGGAGATATACCATGGCCGCCAAGCCTGCTGAAACCAGCA |
| p1002 | AATTCTGTTTCCTGTGTGAAATTGT |

**Supplementary Methods**

**RhlB-mRuby2 purification from *P. aeruginosa*.** 50 ml culture of exponentially growing *P. aeruginosa* cells expressing an -mRuby2 chromosomally tagged RhlB was harvested by centrifugation at 4500 x g for 10 minutes. Pellet was then resuspended in 3,25 mL of lysis buffer (20 mM Tris (pH 8.0), 250 mM NaCl, 10% v/v glycerol, 1/5 EDTA-free protease inhibitor tablet, 50 µg/mL lysozyme) and sonicated for 6x30s, with 15 seconds break in between. After centrifugation of the samples, 25 µL of RFP-TRAP magnetic beads (Chromotek) previously washed in wash buffer (20 mM Tris (pH 8), 250 mM NaCl, 10% v/v glycerol) were added. Incubation on a rotary wheel was done for 1h at 4°C. After that, three wash step with 500 µL of buffer was also performed. Beads were finally covered with 10 µL of sterile PBS and stored at 4°C before LC-MS/MS analysis at the Proteomics Core Facility (University of Geneva). Negative control with untagged wild-type *P. aeruginosa* was analysed in parallel.

**Phylogenetic analyses.** RhlB homologs were identified from previously published analyses (9,10). Amino acid sequences retrieved from the NCBI RefSeq database (11) and aligned with MUSCLE (12). Sequence of *Saccharomyces cerevisiae* EIF4a (ScEIF4a, A6ZQJ1) was added as outgroup. The 983 sequences, together with ScEIF4a, were clustered using *cd-hit* (13) with a 0.9 identity threshold. The representative sequences of the clusters were aligned with *muscle* 3.8.31 (12); the alignment was then refined using the same software. A tree was produced from the alignment using *fasttree* 2.1.10 (14) with the -gamma option and Whelan-And-Goldman 2001 model of amino acid evolution; branches were collapsed below a support value of 0.6 using *TreeGraph* 2.15 (15). The final tree figure was produced thanks to *FigTree* 1.4.4 (http://tree.bio.ed.ac.uk/software/figtree/).

**Protein-protein interaction assays.** The RhlB-RNase E interaction assays were performed exactly as described in the material and methods of Geslain et al., 2025. Briefly, Purified RNase $E^{572\text{-}1057}$- Flag fusion proteins (25 µg) were diluted in buffer H (50 mM Tris-HCl, pH 8.0, 150 mM NaCl, 0.01% Triton X-100) to a final volume of 100 µL and incubated with the magnetic beads for 1 hour at 4 °C on a rotating device. After the incubation, the beads were collected using a magnetic separator and washed three times with 0.25 ml of binding buffer H to remove any unbound protein. the anti-Flag magnetic beads (Sigma-Aldrich) with bound RNase E-Flag fusion proteins, or unbound beads, were incubated with either 5 µg of RhlB or 5 µg of $RhlB^{111\text{-}507}$ diluted in buffer H (final volume 100 µL) for 1 hour at 4 °C on a rotating device. After incubation, the beads were collected using the magnet, and the supernatant (free fraction F) was withdrawn. The anti-Flag beads were then resuspended in 100 µl of binding buffer H and subjected to three rounds of washing of 100 µl each. Following the final wash, the proteins bound to the beads were eluted with 100 µl of 2X LDS sample buffer (Invitrogen) supplemented with 167 mM DTT.

**Western blot analysis.** For each strain, cells density was normalized to an OD600 of 1.0 before loading it onto a 4-12% Bis-Tris mPAGE gel. The gel was migrated and transferred to a nitrocellulose membrane at 3 mA/cm2. After transfer, membranes were blocked overnight in blocking buffer (5% milk powder, 0.1% Tween 20 in Tris-buffered saline, pH 8.0). A 1:10000 dilution of primary anti-Flag antibody (Sigma, F3165) was used. Secondary anti-mouse antibodies conjugated to horseradish peroxidase (Sigma, A9044) were used at a dilution of 1:5000. Western blots were developed using Super-Signal West Pico Chemiluminescent Substrate (Pierce) and visualized on a LAS3000 Fuji Imager.

**Supplementary Figures Legends**

**Figure S1: Phylogenetic analysis of RhlB homologs.** The phylogenetic tress is based on 983 representative amino acid sequences of bacterial RhlB DEAD-box helicases. The *Saccharomyces cerevisiae* protein eIF4a (ScEIF4a) was used to root the tree. Two big clades can readily be identified: Type I RhlB clade including *E. coli* RhlB is colored in black, Type II RhlB clade is highlighted in red and *P. aeruginosa* RhlB sub-group of Type II RhlBs is in yellow.

**Figure S2: Sequence alignment of EcRhlB and PaRhlB proteins using the EBLOSUM62 matrix.** Strictly conserved residues are shown in white on a black background, while partially conserved residues are boxed in grey. Conserved motifs characteristic of DEAD-box RNA helicases (namely Q, I, Ia, Ib, Ic, II, III, IV, IVa, V, Va, and VI) are indicated by grey boxes. The numbers shown above the alignment correspond to the PaRhlB sequence. HDX changes in PaRhlB upon interaction with PaRNase $E^{C\text{-}ter}$ (this study) are highlighted in dark blue, whereas HDX changes in *E. coli* RhlB (EcRhlB) induced by *E. coli* RNase $E^{588\text{-}1061}$ are shown in sky blue (from the HDX-MS results of Bruce et al., 2018).

**Figure S3: Protein purification** Aliquots (2.5 µg) of the eluate fraction of HS-RhlB, HS-$RhlB^{K69A}$ , HS-$RhlB^{1\text{-}111}$, $RhlB^{111\text{-}507}$, RhlB, $RhlB^{1\text{-}111}$, $RhlB^{111\text{-}507}$, HS-RhlB-msfGFP, HS-$RhlB^{111\text{-}507}$-msfGFP, HS-EcRhlB, HS-EcRNase $E^{588\text{-}1061}$, PaRNase $E^{572\text{-}1057}$AR4-REERmut, and PaRNase $E^{572\text{-}1057}$AR1-REERmut were analyzed by SDS-PAGE. The polypeptides were visualized by staining with Coomassie Blue dye. The positions and sizes (kDa) of marker polypeptides are indicated on the left. See Materials and Methods for details on protein purification. Note that PaRNase $E^{572\text{–}1057}$ WT and mutated variants display aberrant migration, as previously reported (see Geslain et al., 2025).

**Figure S4: FRAP experiment.** Top panel: Time-lapse images of in vitro FRAP experiments. The FRAP experiments were performed PaRhlB-msfGFP droplets at 1.75 µM. Bottom panel: Traces of the FRAP data represent mean ± standard deviation from three independent experiments. Half-time and mobile fractions were calculated from the curve.

**Figure S5: The His-Smt3 Tag does not affect RhlB activity, RNA-binding affinity nor LLPS of the purified proteins.** (**A**) In vitro droplet formation of untagged $RhlB^{1\text{-}507}$ and untagged $RhlB^{111\text{-}507}$ at 5 µM or 2.5 µM in the presence of 250 ng/µl poly(U) RNA. Images were acquired using a Nikon Ts2R-FL microscope (40X magnification). Note that no msfGFP-tagged versions were added in this experiment. (**B**) ATPase activity of $His_{10}$Smt3-tagged RhlB, $His_{10}$Smt3-tagged RhlB-K169A, and ULP1-cleaved RhlB. Reaction mixtures (15 µl) containing 250 ng/µl poly(U) RNA (or no RNA, shown as empty symbols) and the indicated enzymes were incubated for 15 minutes at 37 °C. Inorganic phosphate (Pi) release was quantified as described in Materials and Methods and plotted as a function of protein concentration. Data represent the mean of three independent experiments; error bars indicate standard deviation (SD). (**C**) RNA unwinding activity of $His_{10}Smt3$-tagged RhlB and ULP1-cleaved RhlB. Reaction mixtures (15 µl) contained 50 nM 5′-overhang duplex RNA (5′ovh_dsRNA26/14), 3 µM competitor RNA, and 50 nM of the indicated enzymes. The extent of RNA unwinding was measured over time as described in Materials and Methods. (**D**) ATP-dependent RNA binding by $His_{10}Smt3$-tagged RhlB and ULP1-cleaved RhlB. Reaction mixtures (20 µl) contained 1 mM AMP-PNP, 1 nM 5′-FAM-labeled RNA duplex with a 5′ overhang (5′FAM-5′ovh_$dsRNA_{26/14}$) and increasing concentrations of either $His_{10}Smt3$-tagged or ULP1-cleaved RhlB. Samples were incubated at 25 °C for 15 minutes. mFP was measured as described in Materials and Methods and plotted as a function of protein concentration. Data represent the mean of six independent experiments; error bars indicate standard deviation**.** Note that the results for His–Smt3–tagged RhlB shown in panels B, C, and D are identical to those presented in Fig. 2A, 2B, and 2C.

**Figure S6: RNA unwinding activity of *Pa*RhlB.** (**A**) RNA unwinding activity of WT RhlB, and mutant K169A. Reaction mixtures (15 µl) containing 50 nM 5’ overhang duplex RNA (5’ovh_$dsRNA_{26\ /14}$), 3 µM competitor RNA 1 mM ATP and protein as specified. The extent of RNA unwound was determined as described in Materials and Methods and was plotted as a function of time. (**B**) ATP dependence of PaRhlB RNA unwinding. Reaction mixtures (15 µl) containing 50 nM 5’ overhang RNA duplex (5’ovh_$dsRNA_{26/14}$), 3 µM competitor RNA, 1 mM ATP (or no RNA) and enzyme as specified. The extent of RNA unwound was determined as described in Materials and Methods and was plotted as a function of time. (**C**) RNA unwinding specificity. Reaction mixtures (15 µl) containing 50 nM of either 5’ overhang RNA duplex RNA (5’ovh_$dsRNA_{26/14}$), 3’ overhang RNA duplex (3’ovh_$dsRNA_{26\ /14}$), or blunt RNA duplex ($dsRNA_{14/14}$), 3 µM competitor RNA, 1 mM ATP and enzyme as specified. The extent of RNA unwound was determined as described in Materials and Methods and was plotted as a function of time.

**Figure S7: $RhlB^{111\text{-}507}$ directly interacts with PaRNase $E^{572\text{-}1057}$.** (**A**) Aliquots (15 µl) of the input protein sample HS-RhlB full length or 111-507, representing the load (L), of the unbound fraction (U), representing the free fraction, and the bead-bound fraction were mixed with 5 µl of 4X LDS sample buffer (Invitrogen), heated at 80 °C for 5 minutes, and analysed by SDS-PAGE. Polypeptides were visualized using Coomassie Blue staining. (**B**) The control sample showing that PaRhlB and $PaRhlB^{111\text{-}507}$ alone do not bind to the beads are shown.

**Figure S8: Heterologous interaction and ATPase activity stimulation.** (**A**) This figure complements the data shown in Fig. 2E–F. ATPase activity of 1 µM $RhlB^{1\text{-}507}$ or $RhlB^{111\text{-}507}$ in the absence of $RNE^{C\text{-}ter}$ or in the presence of equimolar RNE (1× $PaRNE^{C\text{-}ter}$) or twofold excess of $RNE^{C\text{-}ter}$ (2× PaRNase $E^{572\text{-}1057}$), all measured in the presence of poly(U). (**B**) ATPase activity of 1 µM EcRhlB in the absence (–, buffer) or presence (+) of 2 µM EcRNE$^{C\text{-}ter}$ with the indicated RNA substrates (none, poly(U), totRNA, 5′ovh-dsRNA, or ssRNA) in a buffer containing 50 mM KGlu. (**C**) ATPase activity of 2 µM EcRhlB in the absence (–, buffer) or presence (+) of 4 µM EcRNE$^{C\text{-}ter}$ with the indicated RNA substrates (none, poly(U), totRNA, 5′ovh-dsRNA, or ssRNA) in a buffer containing or 100 mM KCl. Data represent the mean ± standard deviation of three independent experiments.

**Figure S9: HDX-MS mapping of RNase E$^{C\text{-}ter}$ interaction Sites on PaRhlB.** (**A**) Coverage map of PaRhlB showing peptides that were analysed by HDX-MS. Note that the ORF of RhlB starts at Val$^{11}$. (**B**) Deuterium uptake plots (% deuteration ± standard deviation) over time for a representative selection of RhlB peptides, including peptides overlapping RecA1 and RecA2, measured in the presence and absence of PaRNE$^{C\text{-}ter}$. Residue numbers are indicated above each graph. The percentage of maximal deuteration is shown based on the theoretical maximal deuterium uptake. Data are presented as mean ± standard deviation for three independent experiments (cs = charge state). Two of these peptides are also shown in Fig. 3B.

**Figure S10: PaRhlB droplet dissolution: EcRNase E vs PaRNaseE**. Droplets were formed using 1.25 µM PaRhlB in the presence of the indicated RNA substrates (MS2, poly(U), or totRNA) for 60 minutes. After this incubation, either buffer A, 1.25 µM EcRNase E$^{C\text{-}ter}$, or 1.25 µM PaRNase E$^{C\text{-}ter}$ was added to the droplets, and images were taken 60 minutes after addition.

**Figure S11: PaRhlB expression levels.** (**A**) Expression levels of Flag-tagged PaRhlB protein variants shown in Fig 7B were compared by Western blot. (**B**) Growth curves of Δ*rhlB* mutant strains expressing increasing amounts of RhlB (IPTG, 0, 0.15, 0.3, 0.6 or 1.2 mM) at 18 °C. See SI Materials and Methods for experimental details.

#### Supplementary references


1. Green, M.R. and Sambrook, J. (2012) Molecular.cloning.¿a.laboratory.manual. 4th ed. Cold Spring Harbor Laboratory Press, Cold Spring Harbor, N.Y.
2. Holloway, B.W., Krishnapillai, V. and Morgan, A.F. (1979) Chromosomal genetics of Pseudomonas. Microbiol.Rev, **43**, 73-102.
3. Mossessova, E. and Lima, C.D. (2000) Ulp1-SUMO crystal structure and genetic analysis reveal conserved interactions and a regulatory element essential for cell growth in yeast. Mol.Cell, **5**, 865-876.
4. Karimova, G., Ullmann, A. and Ladant, D. (2000) Bordetella.pertussis.adenylate cyclase toxin as a tool to analyze molecular interactions in a bacterial two-hybrid system. Int.J. Med.Microbiol, **290**, 441-445.
5. Voisard, C., Bull, C.T., Keel, C., Laville, J., Maurhofer, M., Schnider, U., Défago, G. and Haas, D. (1994), Molecular.Ecology.of.Rhizosphere.Microorganisms, pp. 67-89.
6. Silva-Rocha, R., Martínez-García, E., Calles, B., Chavarría, M., Arce-Rodríguez, A., de Las Heras, A., Páez-Espino, A.D., Durante-Rodríguez, G., Kim, J., Nikel, P.I..et.al¡ (2013) The Standard European Vector Architecture (SEVA): a coherent platform for the analysis and deployment of complex prokaryotic phenotypes. Nucleic.Acids.Res, **41**, D666-675.
7. Heeb, S., Blumer, C. and Haas, D. (2002) Regulatory RNA as mediator in GacA/RsmA-dependent global control of exoproduct formation in Pseudomonas.fluorescens.CHA0. J. Bacteriol, **184**, 1046-1056.
8. Geslain, S.A.M., Hausmann, S., Geiser, J., Allen, G.E., Gonzalez, D. and Valentini, M. (2025) Critical functions and key interactions mediated by the RNase E scaffolding domain in Pseudomonas aeruginosa. PLoS.genetics, **21**, e1011618.
9. Lopez-Ramirez, V., Alcaraz, L.D., Moreno-Hagelsieb, G. and Olmedo-Alvarez, G. (2011) Phylogenetic distribution and evolutionary history of bacterial DEAD-Box proteins. J.Mol. Evol, **72**, 413-431.
10. Redder, P., Hausmann, S., Khemici, V., Yasrebi, H. and Linder, P. (2015) Bacterial versatility requires DEAD-box RNA helicases. FEMS.microbiology.reviews, **39**, 392-412.
11. Pruitt, K.D., Brown, G.R., Hiatt, S.M., Thibaud-Nissen, F., Astashyn, A., Ermolaeva, O., Farrell, C.M., Hart, J., Landrum, M.J., McGarvey, K.M..et.al¡ (2014) RefSeq: an update on mammalian reference sequences. Nucleic.Acids.Res, **42**, D756-763.
12. Edgar, R.C. (2004) MUSCLE: multiple sequence alignment with high accuracy and high throughput. Nucleic.Acids.Res, **32**, 1792-1797.
13. Li, W. and Godzik, A. (2006) Cd-hit: a fast program for clustering and comparing large sets of protein or nucleotide sequences. Bioinformatics, **22**, 1658-1659.
14. Price, M.N., Dehal, P.S. and Arkin, A.P. (2010) FastTree 2 – Approximately Maximum-Likelihood Trees for Large Alignments. PLOS.ONE, **5**, e9490.
15. Stöver, B.C. and Müller, K.F. (2010) TreeGraph 2: Combining and visualizing evidence from different phylogenetic analyses. BMC.Bioinformatics, **11**, 7.

Fig S1

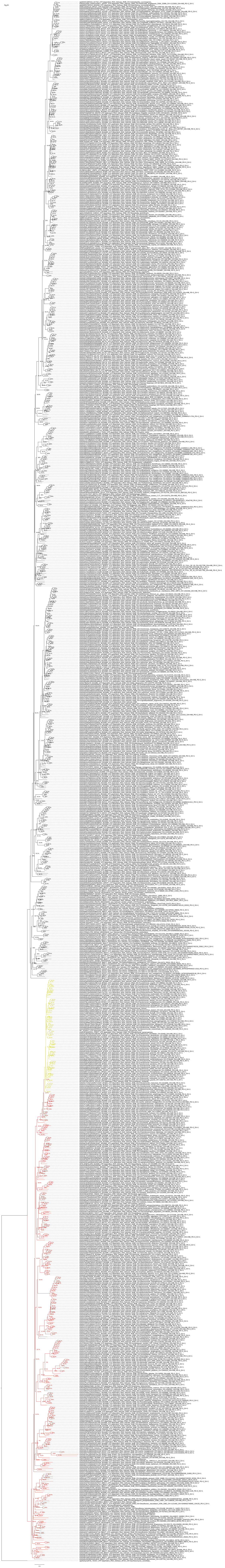

### Figure S2

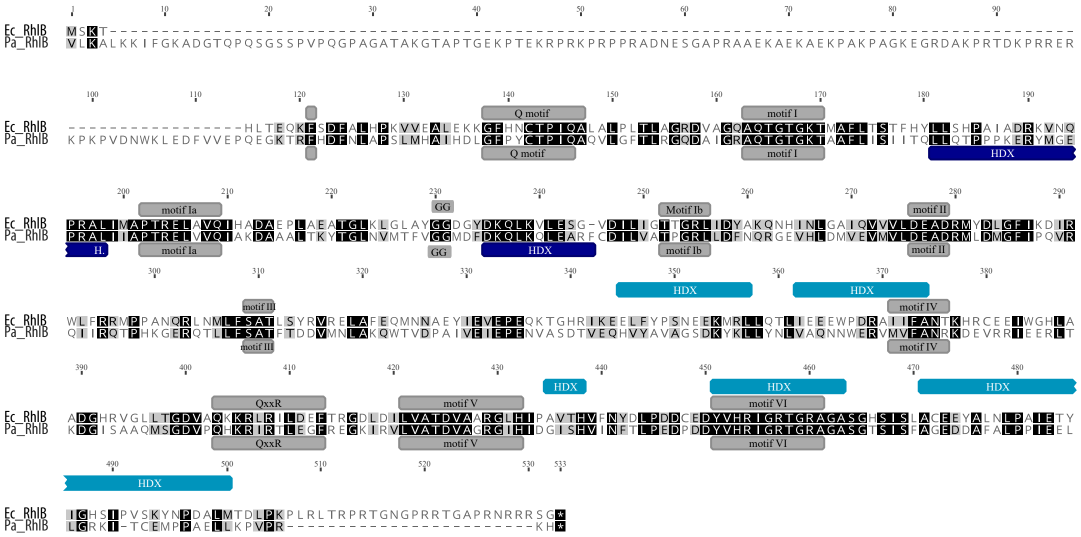
Ec_RhlB
Pa_RhlB
Q motif
motif I
motif Ia
GG
Motif Ib
motif Ib
motif II
motif III
motif IV
QxxR
motif V
motif VI
HDX

Figure S3

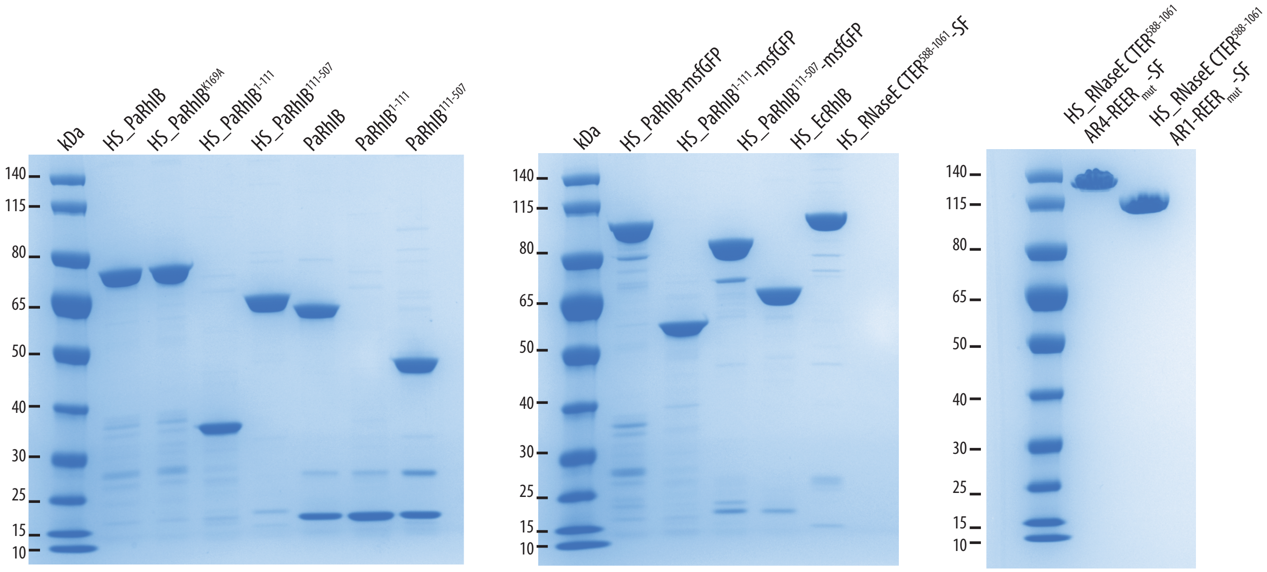

kDa
HS_PaRhlB
HS_PaRhlB$^{K169A}$
HS_PaRhlB$^{1-111}$
HS_PaRhlB$^{111-507}$
PaRhlB
PaRhlB$^{1-111}$
PaRhlB$^{111-507}$
140
115
80
65
50
40
30
25
15
10
kDa
HS_PaRhlB-msfGFP
HS_PaRhlB$^{1-111}$-msfGFP
HS_PaRhlB$^{111-507}$-msfGFP
HS_EcRhlB
HS_RNaseE CTER$^{588-1061}$-SF
HS_RNaseE CTER$^{588-1061}$ AR4-REER$_{mut}$-SF
HS_RNaseE CTER$^{588-1061}$ AR1-REER$_{mut}$-SF

Figure S4

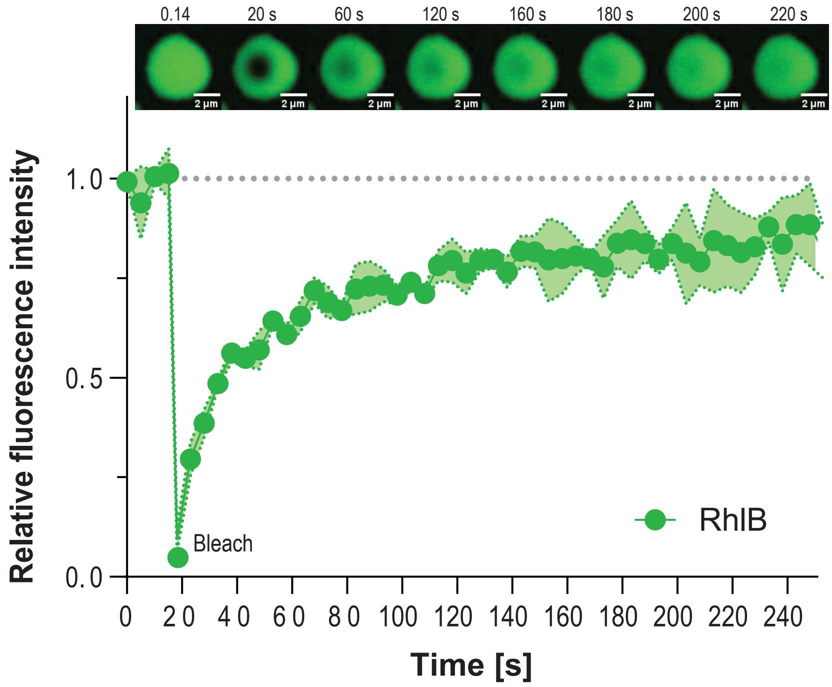
0.14
20 s
60 s
120 s
160 s
180 s
200 s
220 s
2 µm
Relative fluorescence intensity
1.0
0.5
0.0
Bleach
RhlB
0
20
40
60
80
100
120
140
160
180
200
220
240
Time [s]

Figure S5

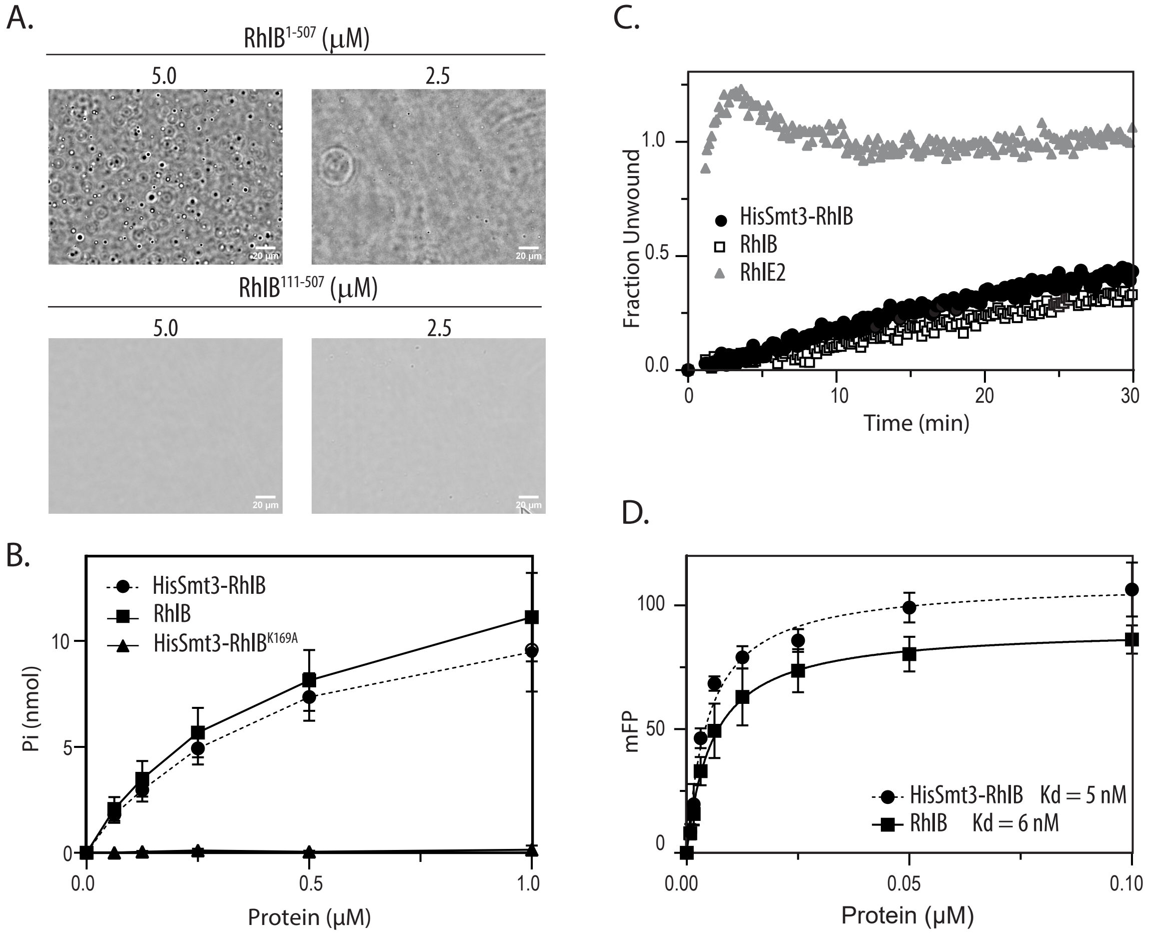
A.
RhlB$^{1-507}$ (μM)
5.0
2.5
RhlB$^{111-507}$ (μM)
5.0
2.5
B.
HisSmt3-RhlB
RhlB
HisSmt3-RhlB$^{K169A}$
Pi (nmol)
Protein (μM)
C.
Fraction Unwound
HisSmt3-RhlB
RhlB
RhlE2
Time (min)
D.
mFP
HisSmt3-RhlB Kd = 5 nM
RhlB Kd = 6 nM
Protein (μM)

Figure S6

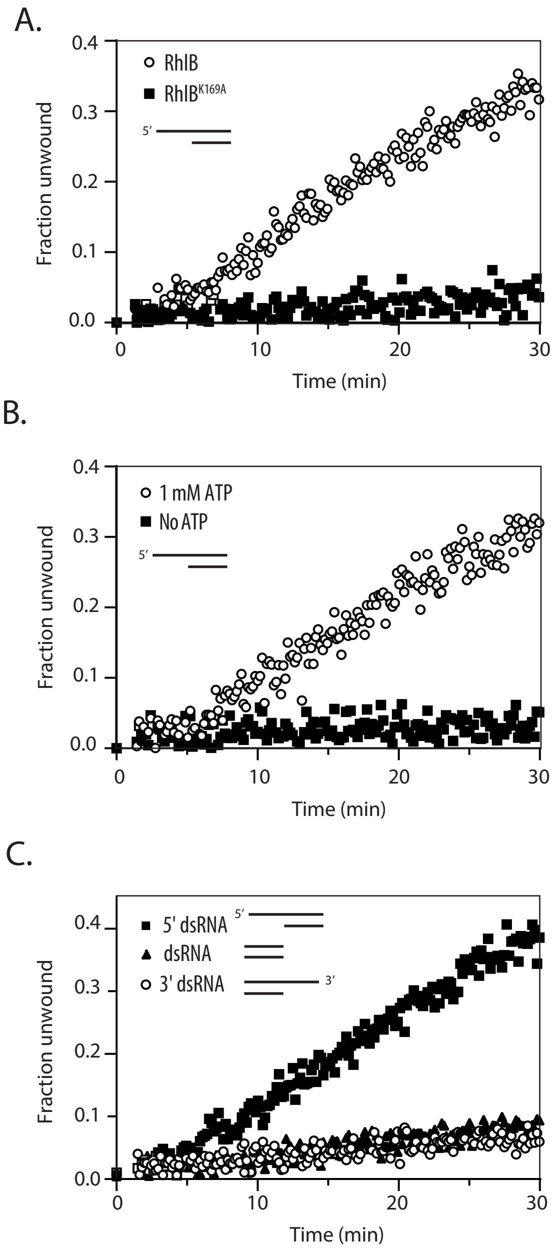
A.
Fraction unwound
Time (min)
RhlB
RhlB K169A
5′
0.0
0.1
0.2
0.3
0.4
0
10
20
30
B.
Fraction unwound
Time (min)
1 mM ATP
No ATP
5′
0.0
0.1
0.2
0.3
0.4
0
10
20
30
C.
Fraction unwound
Time (min)
5' dsRNA
dsRNA
3' dsRNA
5′
3′
0.0
0.1
0.2
0.3
0.4
0
10
20
30

Figure S7

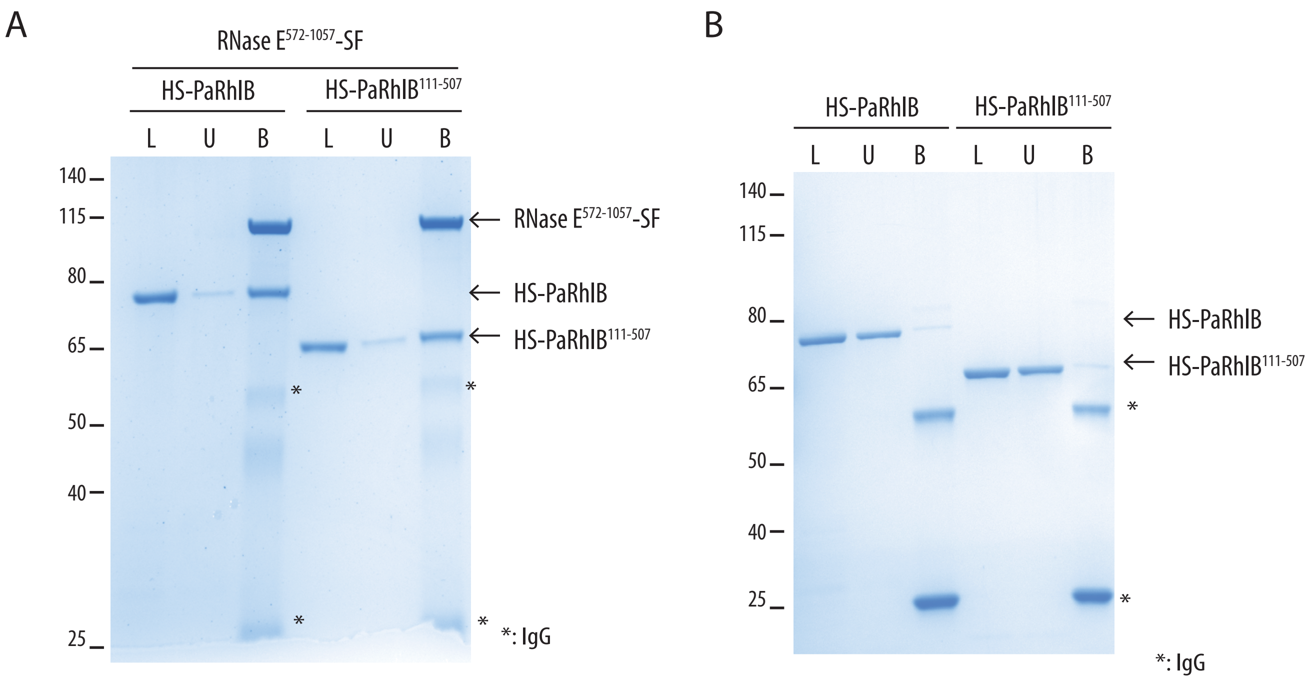
A
RNase E572-1057-SF
HS-PaRhlB
HS-PaRhlB111-507
L U B L U B
140
115
80
65
50
40
25
RNase E572-1057-SF
HS-PaRhlB
HS-PaRhlB111-507
*
*
*
*
*: IgG
B
HS-PaRhlB
HS-PaRhlB111-507
L U B L U B
140
115
80
65
50
40
25
HS-PaRhlB
HS-PaRhlB111-507
*
*
*: IgG

Figure S8

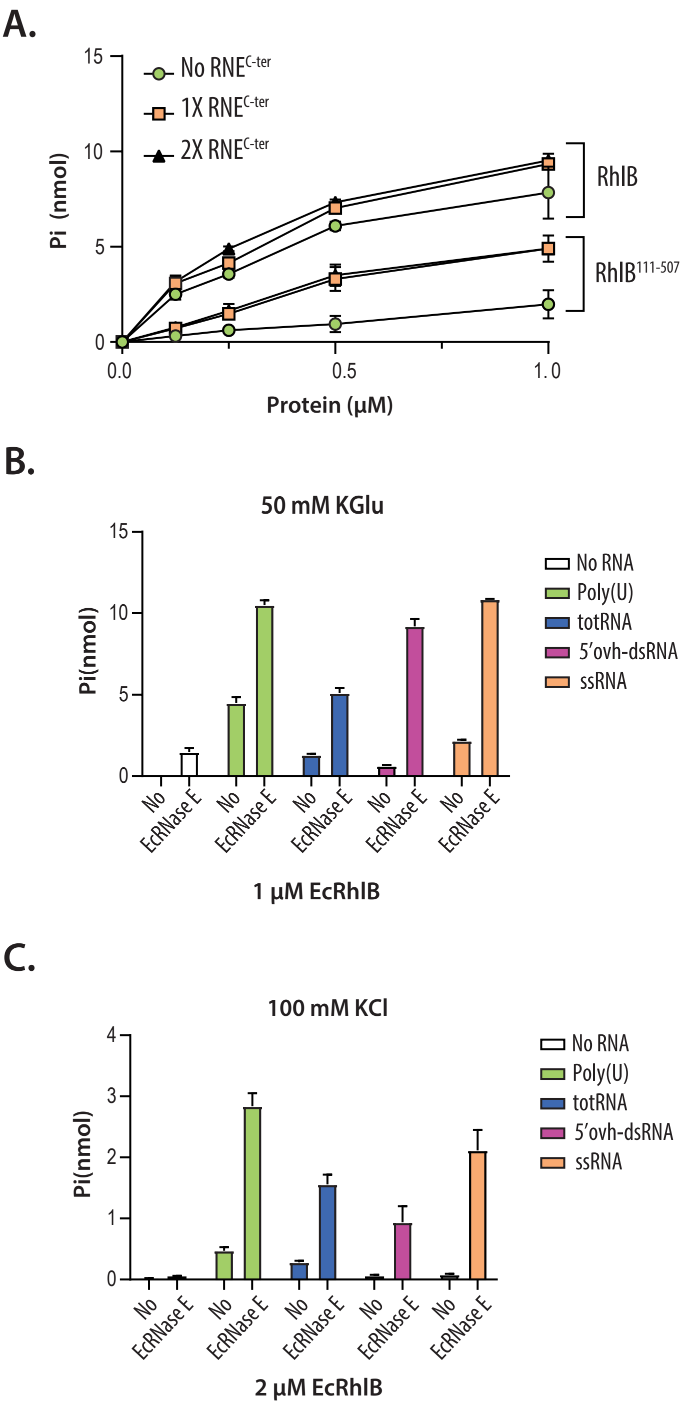

A.
15
10
5
0
Pi (nmol)
No RNE^C-ter
1X RNE^C-ter
2X RNE^C-ter
RhlB
RhlB^111-507
0.0
0.5
1.0
Protein (μM)
B.
50 mM KGlu
15
10
5
0
Pi(nmol)
No RNA
Poly(U)
totRNA
5'ovh-dsRNA
ssRNA
No
EcRNase E
1 μM EcRhlB
C.
100 mM KCl
4
3
2
1
0
Pi(nmol)
No RNA
Poly(U)
totRNA
5'ovh-dsRNA
ssRNA
No
EcRNase E
2 μM EcRhlB

Figure S9

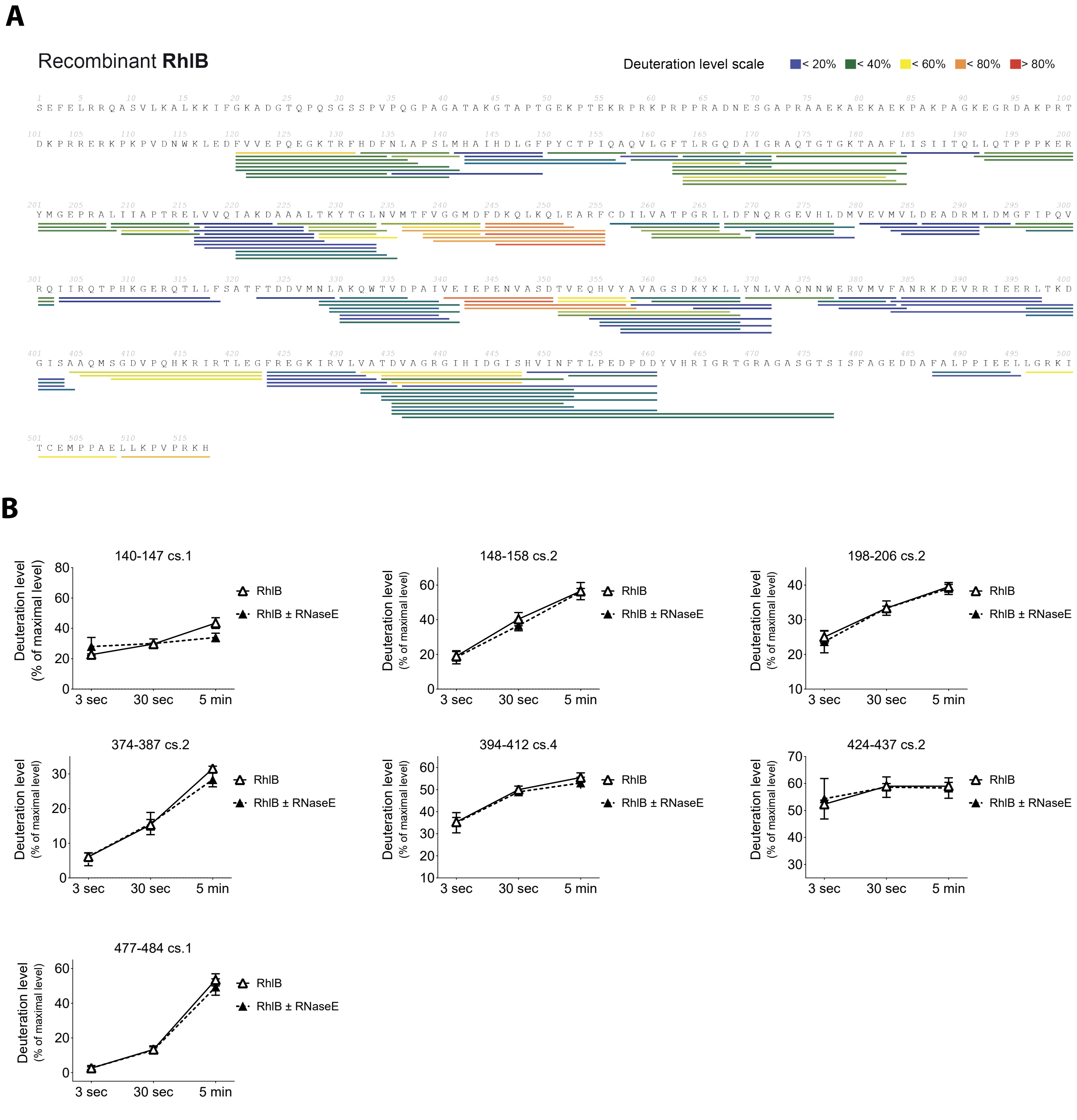
A
Recombinant RhlB
Deuteration level scale
< 20%
< 40%
< 60%
< 80%
> 80%
B
140-147 cs.1
148-158 cs.2
198-206 cs.2
374-387 cs.2
394-412 cs.4
424-437 cs.2
477-484 cs.1
Deuteration level
(% of maximal level)
3 sec
30 sec
5 min
RhlB
RhlB ± RNaseE

Figure S10

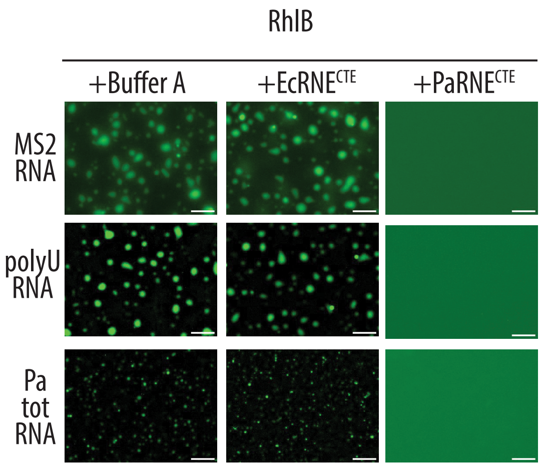

Figure S11

**A.**

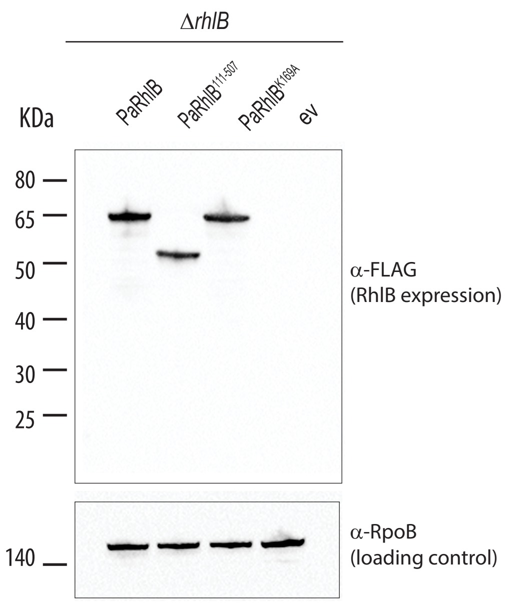


**B.**

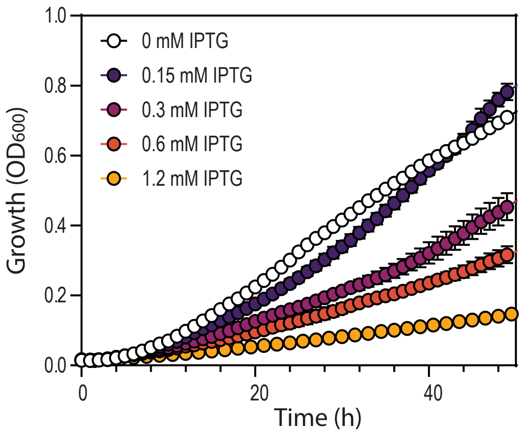
